# Resource supply dynamics control stability and chaos in complex ecosystems

**DOI:** 10.64898/2026.09.01.748502

**Authors:** Jamila Rowland-Chandler, Akshit Goyal, Wenying Shou

## Abstract

Ecological interactions are often mediated by feedbacks between organisms and their resource environments. Yet, how resource supply dynamics dictate collective dynamical phases of an ecosystem remains unclear. Here, we analyse a generalised consumer–resource model with non-reciprocal interactions to demonstrate that self-renewing versus externally-supplied resources yield fundamentally different dynamical phase diagrams. As interactions become increasingly non-reciprocal, ecosystems relying on self-renewing resources transition from stable dynamics to chaos and ultimately to infeasibility. By contrast, ecosystems with externally-supplied resources remain stable over a broader parameter range and transition to infeasibility without experiencing an intervening chaotic phase. Using the cavity method, we derive a unified stability condition applicable to a broad class of resource supply functions, explaining why externally-supplied resources can expand the stable region. We show that stability hinges crucially on the susceptibility of resources to perturbations, which depends strongly on their supply. Further, we show that external resource supply suppresses chaos in the unstable region by drastically reducing the susceptibility of resources closest to extinction. Our findings demonstrate that resource dynamics fundamentally reshape the accessible dynamical behaviours of an ecosystem, with implications for interpreting microbial community experiments.

---

A central goal of ecology is to understand how collective dynamical behaviours in ecosystems, from stable equilibria to chaotic fluctuations [1–10], emerge from interactions among many species. Recent approaches from statistical physics have made significant progress on this problem by treating ecosystems as large disordered dynamical systems, revealing novel dynamical phases and phase transitions controlled by the statistical properties of ecological interactions [5, 11–27].

Much of this work has focused on generalised Lotka– Volterra (gLV) models with direct pairwise species interactions [5, 12–16, 20, 22–26], which lack explicit feedbacks between organisms and their environments that often mediate species interactions [10, 28–44]. This limitation has motivated growing interest in consumer–resource models (CRM) where species interactions emerge from competition and cooperation for shared resources [17– 19, 21, 27, 33, 45–54]. Recent work has shown that decreasing the reciprocity between resource depletion and consumer growth can drive a transition from stability to chaotic abundance fluctuations [17–19, 27]. However, these models assume self-renewing resources; whether these results extend to externally supplied resources remains unclear, despite some studies claim that they do [18, 19]. As a result, we lack a general understanding of how resource supply modes shape the dynamical phases of ecosystems, hindering interpretation of recent experiments linking resource supplies to dynamical fluctuations [5, 6, 55].

In this Letter, we show that changing resource supply mode fundamentally alters the phase diagram of dynamical states in a complex consumer–resource ecosystem. As interaction reciprocity decreases, while ecosystems with self-renewing resources transition from stability to chaotic boom–bust dynamics and ultimately to infeasibility, ecosystems with externally-supplied resources remain stable over a wider parameter range and transition directly to infeasibility. Using the cavity method, we derive a unified stability condition for a broad class of consumer–resource models, and show that different resource supply modes lead to distinct stability conditions depending on whether resources can go extinct. Our results demonstrate that the mode of resource supply can determine which dynamical behaviours an ecosystem can exhibit.

## Model

We consider a generalised consumer-resource model with non-reciprocal interactions, where a pool of *S* consumer species grow from consuming *M* substitutable resources (Fig. 1). The dynamics of resource *α* (*R*_*α*_, *α* ∈ {1, …*M*}) and consumer *i* (*N*_*i*_, *i*∈ {1, …*S*}) are described by the following set of differential equations:

**FIG. 1.**
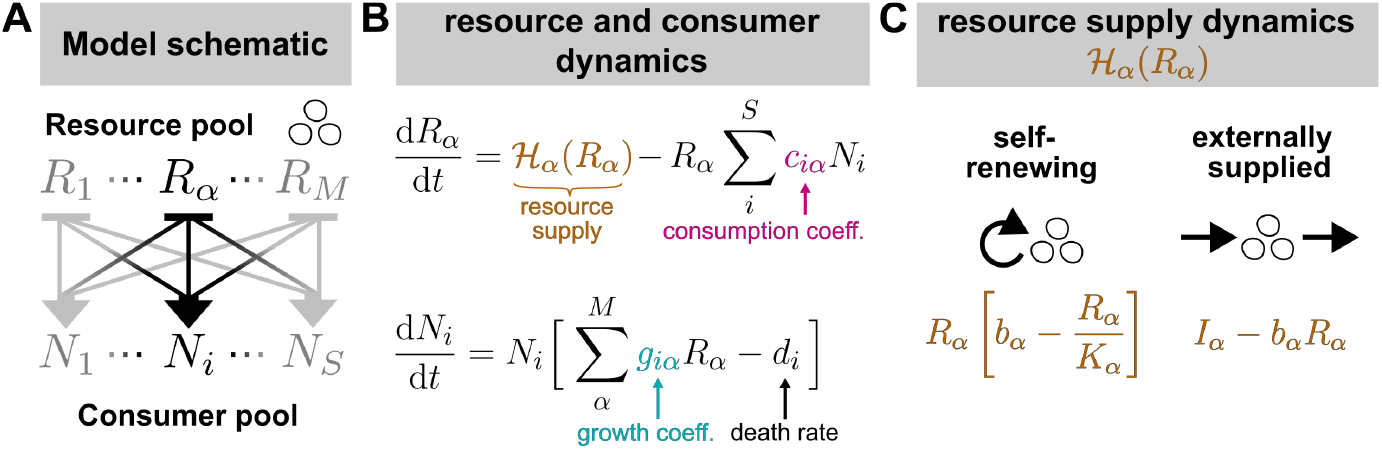
Schematic for generalised consumer-resource models with non-reciprocal interactions. **(A)** Interactions between the resource and consumer pool. **(B)** General form of resource and consumer dynamics. **(C)** Resource supply functions investigated in this paper.

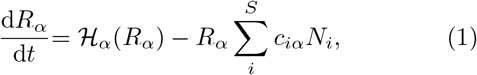

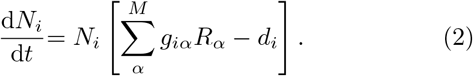

Here, *c*_*iα*_ is the per-capita consumption rate of resource *α* by consumer *i, g*_*iα*_ is the corresponding growth rate of consumer *i* on resource *α*, and *d*_*i*_ is consumer *i*’s death rate. *H*_*α*_(*R*_*α*_) is the resource supply function, which describes resource dynamics in the absence of consumers. When resources are self-renewing or biological (e.g., bacteria are resources for phage), *H*_*α*_(*R*_*α*_) describes logistic growth:

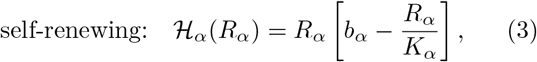

where *b*_*α*_ is the intrinsic growth rate of resource *α* and *K*_*α*_ is its carrying capacity. When resources are externally-supplied or abiotic (e.g., chemicals supplied in a chemostat for bacterial growth), *H*_*α*_(*R*_*α*_) takes the form

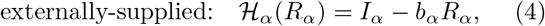

where *I*_*α*_ is the influx rate of resource *α*, independent of resource density, and *b*_*α*_ is now interpreted as the outflux or dilution rate. When resource dynamics are much faster than consumer dynamics, both classes of dynamics reduce to gLV model and can become indistinguishable under certain conditions (SI Appendix E). Here, we investigate the opposite limit where resource environments actively shape species interactions, and resource dynamics cannot be ignored.

## Parametrisation

;To study the typical dynamics of highly-diverse communities (*M*, ≫ *S* 1), following prior literature, we sample model parameters as random variables [12, 14, 24, 48] drawn from fixed distributions:

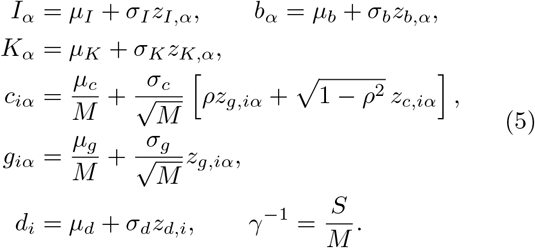

where all *z*_*⟨·⟩*_ terms represent uncorrelated standard Gaussian random variables (note that our results also generalise to non-Gaussian distributions; SI Appendix B). Consumption (*c*_*iα*_) and growth (*g*_*iα*_) rates are scaled by the resource pool size *M* to ensure a proper thermodynamic limit (*M* → ∞, *S* → ∞) while maintaining a finite ratio *γ*^*−*1^. For simplicity, we set *µ*_*c*_ = *µ*_*g*_ ≡ *µ* and *σ*_*c*_ = *σ*_*g*_ ≡ *σ* (quantifying heterogeneity) in all simulations. The parameter *ρ* = corr(*c*_*iα*_, *g*_*iα*_) represents the reciprocity of consumer–resource interactions: when *ρ* = 1, *c*_*iα*_ = *g*_*iα*_, whereas when *ρ* = 0 they are independent.

## Resource supply mode sets accessible dynamical phases

Previous work on consumer-resource models with self-renewing resources showed that the dynamical phase diagram can be plotted in terms of the reciprocity *ρ* and heterogeneity *σ* in consumption and growth [17, 18, 27]. Decreasing *ρ* (or increasing *σ*) drives communities through two successive transitions: from stable to chaotic dynamics, and then to infeasible dynamics where abundances could grow unbounded (Fig. 2A, B); [17]). To test if changing the mode of resource supply impacts stability we simulated both self-renewing and externally-supplied resource models and plotted the dynamical phase diagram, taking care to match consumer and resource statistics in both cases (SI Appendix D).

**FIG. 2.**
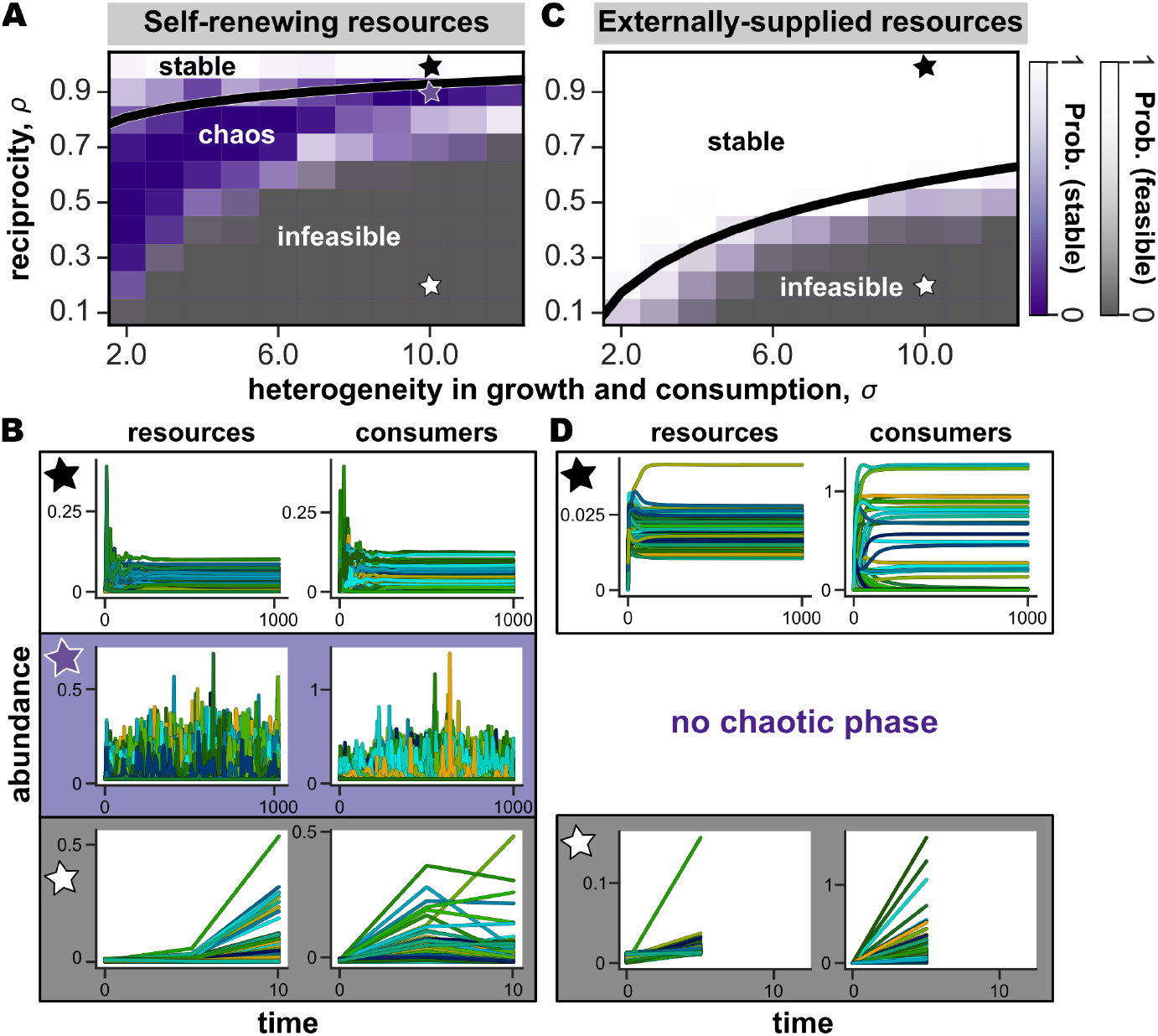
The mode of resource supply governs accessible dynamical phases and the size of stable region. **(A, B)** Consumer-resource model with self-renewing resources. (A) Phase diagram. Each cell shows results from 20 simulated communities, with darker purple indicating a higher proportion of simulations showing unstable but feasible dynamics (maximum Lyapunov exponent *>* 0), darker grey indicating a higher proportion showing infeasible dynamics (simulations are numerically unstable and thus terminate early), and white indicating stable communities. Black line = analytically-derived stability condition *ρ*^2^ = *S*^*∗*^*/M* ^*∗*^, where *S*^*∗*^*/M* ^*∗*^ was estimated from simulations. (B) Representative dynamics from the stable (black star, top), chaotic (purple star, middle) and infeasible regions (white star, bottom). Each curve in each plot represents a single consumer or resource. **(C)** Phase diagram for the model with externally-supplied resources. Black line = analytically-derived stability condition in a simple limit, *ρ*^2^ = *S*^*∗*^*/M*, where *S*^*∗*^*/M* was estimated from simulations. **(D)** Representative dynamics from the stable and infeasible regions of the externally-supplied model. There is no chaotic phase. (Simulation parameters are given in SI Appendix D.)

Surprisingly, the externally-supplied model remained stable over a far greater range of *ρ* and *σ* than the self-renewing model (Fig. 2C). Moreover, when the externally-supplied model did lose stability, it bypassed the chaotic regime entirely and instead transitioned from globally stable directly to infeasible dynamics (Fig 2C, D). Any chaotic trajectories observed near the stability-infeasibility boundary (Fig. 2C, light purple) were likely due to finite size effects, and needed parameters to be extremely fine-tuned to be observed (SI Appendix E Fig. S2). These results demonstrate that ecosystems with externally-supplied resources have qualitatively different collective dynamics than with self-renewing resources.

## Unified stability condition

To understand what drives the difference in dynamical regimes when changing resource supply mode, we used the cavity method to derive a unified stability condition for consumer–resource models with an arbitrary resource supply function *H*_*α*_(*R*_*α*_) (SI Appendices A–B). The cavity method involves deriving self-consistency equations for the steady-state abundance distributions of consumers and resources [14, 17, 20, 48]. When the moments of the distributions corresponding to species and resource responses to perturbations diverge, communities become unstable. Applying this to the general class of consumer–resource models in Eqs. (1)–(2), we obtain the following unified stability condition:

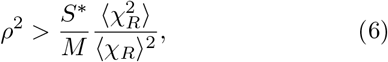

where *S*^*∗*^ is the number of surviving consumers, and *χ*_*R*_ = {−*∂R*_*α*_*/∂b*_*α*_} is the distribution of resource self-susceptibilities describing how a resource’s abundance responds to perturbations in its growth or dilution rate.

To illustrate how this condition manifests for both models, we consider the simple limit of homogeneous resources (*σ*_*b*_ = *σ*_*K*_ = *σ*_*I*_ = 0). In this limit, the key difference between the two models is that self-renewing resources can go extinct (*M* ^*∗*^≤ *M* resources survive), while externally-supplied resources cannot go extinct because of the constant influx *I*_*α*_ ≠ 0. As a result, the ratio 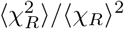 simplifies to *M/M* ^*∗*^ for self-renewing, and 1 for externally-supplied resources (SI Appendix B), and the stability conditions reduce to:

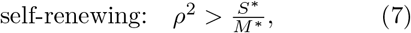

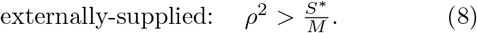

Since ecosystems with externally-supplied resources are typically less niche-packed than self-renewing resources (*S*^*∗*^*/M*≤ 0.5 for externally-supplied, *S*^*∗*^*/M* ^*∗*^≤1 for self-renewing [51]), Eqs. (7)–(8) help explain that externally-supplied resources should typically support a larger stable region. Note that our unified condition recovers the known stability condition for self-renewing resources, Eq. (7) [17, 18].

When resources are heterogeneous, Eq. (8) no longer holds and we must instead obtain 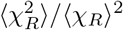 using the self-consistency equations (SI Appendices A–B). This results in a more general stability condition for externally-supplied resources, which is rather different from that for self-renewing resources (Eq. (7)):

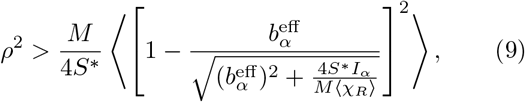

where 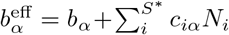 is the effective depletion rate of resource *α* due to outflux and consumption. Eq. (9) agrees remarkably well with our numerical phase diagrams (SI Appendix E Fig. S1 A–B). In sum, our unified condition shows that stability depends not just on the interplay between reciprocity and niche packing [17, 18], but also importantly on resource susceptibilities, specifically whether resources can go extinct. This difference in resource susceptibilities is key to why changing resource supply mode can drastically change the stable region.

## External resource influx suppresses chaos

Once unstable, the externally-supplied resource model does not exhibit chaos. To understand what drives this loss of chaos, we investigated a hybrid model combining the two resource supply functions in Eqs. (3)–(4):

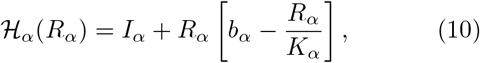

where *I*_*α*_ is the externally-supplied influx rate, *b*_*α*_ can be the intrinsic resource growth rate or dilution rate (depending on its sign), and 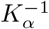 is the strength of resource self-inhibition. The hybrid model could describe, e.g., the resource-consumer dynamics of bacteria–phages in sewers, with bacteria supplied by both self-renewal and external inflow. External influx (*I*_*α*_) is unique to the externally-supplied model, while self-inhibition 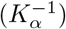 is unique to the self-renewing model. Thus, tuning these terms enables the hybrid model to interpolate between the two supply functions.

We simulated phase diagrams for this model spanning four cases: in the presence or absence of influx and self-inhibition (Fig. 3 A; SI Appendix D for details). Without resource influx, the hybrid model had a small or no stable phase and a large chaotic phase, irrespective of self-limitation (Fig. 3 A, left two panels). On the other hand, with resource influx, the stable phase expanded and chaos was suppressed (Fig. 3 A, right two panels). To quantitatively observe the loss of chaos, we performed additional simulations in which we systematically increased the influx rate. Doing so systematically broadened the stable phase and suppressed both chaotic and infeasible phases, but the chaotic phase was suppressed faster, disappearing entirely at *I*_*α*_ ≥10^*−*2^ (Fig. 3 B). Analytic cavity calculations of the stability condition for the hybrid model were consistent with our unified stability condition Eq. (6) (SI Appendix C). When evaluated explicitly, this condition resembled that of the externally-supplied model (SI Appendix C). This shows that external resource influx is a stronger driver of phase transitions and chaos than resource self-inhibition.

**FIG. 3.**
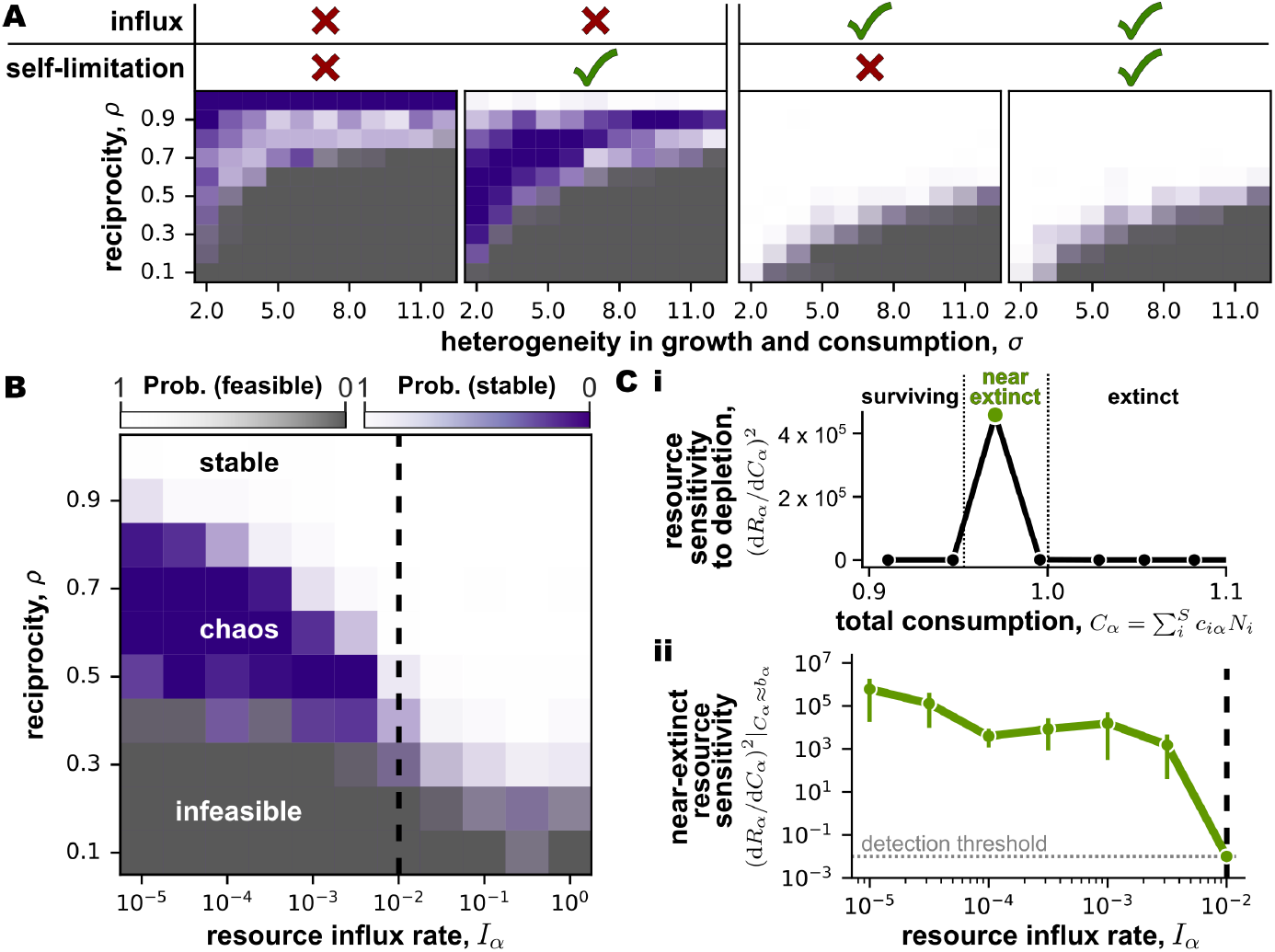
External influx of resources suppresses chaos. (Simulation parameters are given in SI Appendix D.) **(A)** Phase diagrams of the hybrid model, where resources are both externally-supplied and self-renewing: *H*_*α*_(*R*_*α*_) = *I*_*α*_ + *R*_*α*_ [*b*_*α*_*− R*_*α*_*/K*_*α*_]. Approximate presence and absence of resource influx *I*_*α*_ and self-inhibition 1*/K*_*α*_ is indicated by a tick or cross. Refer to Fig. 2 for details on how the diagrams was generated. **(B)** Phase diagram for the hybrid model as a function of reciprocity and external resource influx *I*_*α*_, self-inhibition 1*/K*_*α*_ = 1. **(C)** (i) Example binned distribution of resource sensitivities to changes in their total depletion rate *C*_*α*_, sampled at *I*_*α*_ = 10^*−*5^, number of bins = 15. The green point is the sensitivity of near extinction resources (*C*_*α*_ *≈b*_*α*_*≈* 1) used in plot C ii. (ii) Sensitivity of near-extinct resources to consumption (similar to (i)) as a function of influx rate along the stability-chaos boundary (Appendix D for details). The detection threshold for peaks in sensitivity was set to 0.01 (dashed grey vertical line), so at *I*_*α*_ = 10^*−*2^ there was no detectable peak. Points indicate average over realizations, error bars are 95% confidence intervals for *n* = 80 communities.

We now explain how increasing influx leads to loss of chaos. In consumer-resource models, a resource is driven towards its baseline abundance when it is heavily consumed (with total consumption 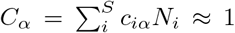) with the baseline set by the resource influx rate. Thus, when there is no influx the baseline is zero, so resources can go extinct. In the chaotic phase, near-baseline resources are highly sensitive to changes in their total depletion, which is typically quantified by the second moment of the response [23], and in this case given by 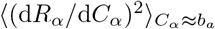(Fig. 3 C i).

One might naïvely expect that raising influx will not affect chaos, since increasing the baseline resource abundance supports larger consumer populations and hence greater depletion. However, as the rate of resource influx increases, resource concentrations become controlled more by influx than by consumption. Consequently, resources that are close to extinction may no longer be highly sensitive to perturbations such as consumer depletion. To test this, we varied resource influx in the hybrid model and measured the sensitivity of near-extinction resources 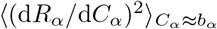 near the stability boundary (see SI Appendix D for details). We found that increasing resource influx decreased the sensitivity of near-extinct resources by several orders of magnitude. Sensitivity became practically non-detectable (≪1) at *I*_*α*_ ≥10^*−*2^ (Fig. 3 C ii), exactly the influx rate at which we observed the loss of the chaotic phase (Fig. 3 B). This confirmed that external resource influx suppresses chaos by reducing the sensitivity of resources to consumer depletion.

## Discussion

Our central finding is that the mode of resource supply can fundamentally alter the dynamical collective phases of an ecosystem. While self-renewing resources yield stable, chaotic, and infeasible phases, externally-supplied resources yield only stable and infeasible phases, without chaos. The unified stability condition shows that this difference arises because resources can go extinct under self-renewing but not external resource supply. External supply then suppresses chaos by desensitising near-extinction resources to perturbations.

Our results contrast with previous work suggesting that resource supply does not alter ecosystem phase transitions [18, 19]. Those studies prescribed fixed points with a chosen set of surviving species and resources, rather than letting them emerge from ecological dynamics. As a result, they likely also implicitly set the resource susceptibilities to be the same for both forms of resource supply. Thus it might appear that resource supply has no effect on ecosystem stability. Further work is needed to establish the source of the observed discrepancy.

Recent experiments show microbial communities exhibiting persistent fluctuations under external resource supply [4, 5, 55], yet our theory predicts suppression of chaos. This suggests that the fluctuations observed in these experiments might arise from mechanisms other than resource-mediated interactions. Examples include pH modification [6], direct interactions [5, 15, 20, 23, 56, 57], and periodic resource supply (serial dilution) in experiments versus constant resource supply (chemostats). Identifying which of these mechanisms underlie chaos is key to controlling dynamical phases in natural and laboratory communities.

Finally, our results highlight that resource supply mode can fundamentally alter feasible phases of ecosystem dynamics. Our unified stability condition could help classify other modes of resource supply, e.g., cross-feeding, by the dynamical phases they support (SI Appendix B). Analysing the chaotic phase using dynamical mean-field theory [20, 23, 58] would also shed further light on the quantitative patterns of dynamical phases under different resource supply modes. These directions would lead to a better understanding of different dynamical behaviours observed in natural ecosystems.

## Acknowledgements

We thank G. Bunin, R. Mallikarjun, S. Pollak, P. Mehta, and E. Blumenthal for valuable discussions. This research was supported in part by grant NSF PHY-2309135 to the Kavli Institute for Theoretical Physics (KITP) and the Gordon and Betty Moore Foundation Grant No. 2019.02. A.G. acknowledges support from the DAE under project no. RTI4001, an Ashok and Gita Vaish Junior Researcher Award, as well as the Centre for Artificial Learning and Intelligence for Biological Research and Education, ICTS-TIFR. W.S. acknowledges support from the Royal Society Wolfson Fellowship, as well as an Academy of Medical Sciences Professorship.

