## Supplementary Materials for "Resource supply dynamics control stability and chaos in complex ecosystems"

### Supplementary Information for "Resource supply dynamics control stability and chaos in complex ecosystems"

### A Stability condition for externally-supplied resources using the cavity method

#### A.1 Steady-state community before introducing the cavity consumer and resource

We begin with a steady-state community containing $S$ consumers and $M$ resources before the cavity consumer and resource 0 are introduced. The steady-state abundances of resident consumers and resources are described by the following equations:

$$\frac{\mathrm{d}N_{i\setminus 0}}{\mathrm{d}t} = 0 = N_{i\setminus 0}\left[\sum_{\alpha=1}^{M} g_{i\alpha}R_{\alpha\setminus 0} - d_i\right]. \tag{1}$$

$$\frac{\mathrm{d}R_{\alpha\setminus 0}}{\mathrm{d}t} = 0 = I_\alpha - b_\alpha R_{\alpha\setminus 0} - R_{\alpha\setminus 0}\sum_{i=1}^{S} c_{i\alpha}N_{i\setminus 0}. \tag{2}$$

Next, we substitute the growth and consumption coefficients with their parameter sampling rules (Main text, Parametrisation). Starting with the resident consumer, we obtain

$$0 = N_{i\setminus 0}\left[\sum_{\alpha=1}^{M}\left[\frac{\mu_g}{M} + \frac{\sigma_g}{\sqrt{M}}z_{g,i\alpha}\right]R_{\alpha\setminus 0} - d_i\right] = N_{i\setminus 0}\left[\mu_g\langle R\rangle + \frac{\sigma_g}{\sqrt{M}}\sum_{\alpha=1}^{M} z_{g,i\alpha}R_{\alpha\setminus 0} - d_i\right]. \tag{3}$$

Note that $\mu_g\langle R\rangle = [\mu_g/M]\sum_{\alpha=1}^{M} R_{\alpha\setminus 0}$, where $\langle R\rangle$ is the average abundance of resident resources, is an expression we define self-consistently later.

Repeating the same process with resident resources gives us

$$0 = I_\alpha - b_\alpha R_{\alpha\setminus 0} - R_{\alpha\setminus 0}\left[\mu_c\gamma^{-1}\langle N\rangle + \frac{\sigma_c}{\sqrt{M}}\sum_{i=1}^{S}\left[\rho z_{g,i\alpha} + \sqrt{1-\rho^2}z_{c,i\alpha}\right]N_{i\setminus 0}.\right]. \tag{4}$$

Note that $\mu_c\gamma^{-1}\langle N\rangle = [\mu_c/M]\sum_{i=1}^{S} N_{i\setminus 0}$, where $\langle N\rangle$ is the average abundance of resident consumers, is an expression we define self-consistently later.

#### A.2 Introducing the cavity consumer and resource

Next, we invade the existing community with the cavity consumer and resource, which are statistically identical to the resident consumers and resources. The steady-state abundances for the resident consumers after invasion $N_i$ and resources $R_\alpha$ thus obey the following equations:

$$\begin{aligned}\frac{\mathrm{d}N_i}{\mathrm{d}t} = 0 &= N_i\Big[\sum_{\alpha=1}^{M} g_{i\alpha}R_\alpha - d_i + g_{i0}R_0\Big]\\ &= N_i\Big[\mu_g\langle R\rangle + \frac{\sigma_g}{\sqrt{M}}\sum_{\alpha=1}^{M} z_{g,i\alpha}R_\alpha - d_i + \frac{\sigma_g}{\sqrt{M}}z_{g,i0}R_0\Big].\end{aligned} \tag{5}$$

The effect of the cavity resource on the dynamics of consumer $i$ — $\sigma_g/\sqrt{M}\sum_{\alpha=1}^{M} z_{g,i0}R_0$ — can be interpreted as a small perturbation to the death rate of consumer $i$, $\Delta d_i$:

$$d_i \xrightarrow{\text{perturb}} d_i - \frac{\sigma_g}{\sqrt{M}} z_{g,i0} R_0. \tag{6}$$

Similarly, the new steady-state abundances of resident resources after invasion by the cavity variables obey the following equations:

$$\frac{\mathrm{d}R_\alpha}{\mathrm{d}t} = 0 = I_\alpha - b_\alpha R_\alpha - R_\alpha \left[\mu_c \gamma^{-1}\langle N\rangle + \frac{\sigma_c}{\sqrt{M}} \sum_{i=1}^{S} \left[\rho z_{g,i\alpha} + \sqrt{1-\rho^2} z_{c,i\alpha}\right] N_i\right] - \frac{\sigma_c}{\sqrt{M}} R_\alpha \left[\rho z_{g,0\alpha} + \sqrt{1-\rho^2} z_{c,0\alpha}\right] N_0. \tag{7}$$

This effect of the cavity consumer on the dynamics of resource $\alpha$ can be interpreted as a small perturbation to the outflux rate of resource $\alpha$, $\Delta b_\alpha$:

$$b_\alpha \xrightarrow{\text{perturb}} b_\alpha + \frac{\sigma_c}{\sqrt{M}} \left[\rho z_{g,0\alpha} + \sqrt{1-\rho^2} z_{c,0\alpha}\right] N_0. \tag{8}$$

We can relate the post-perturbation abundance of consumer $i$, $N_i$ with its pre-perturbation abundance $N_{i\backslash 0}$ using a linear response approximation, given by:

$$\begin{aligned} N_i &\approx N_{i\backslash 0} + \sum_{j=1}^{S} \underbrace{\frac{\partial N_i}{\partial d_j}}_{v_{N,ij}} \Delta d_j + \sum_{\beta=1}^{M} \underbrace{\frac{\partial N_i}{\partial b_\beta}}_{-\chi_{N,i\beta}} \Delta b_\beta. \\ &\approx N_{i\backslash 0} - \sum_{j=1}^{S} v_{N,ij} \left[\frac{\sigma_g}{\sqrt{M}} z_{g,j0} R_0\right] - \sum_{\beta=1}^{M} \chi_{N,i\beta} \left[\frac{\sigma_c}{\sqrt{M}} \left[\rho z_{g,0\beta} + \sqrt{1-\rho^2} z_{c,0\beta}\right] N_0\right], \\ &\text{where } v_{N,ij} = \frac{\partial N_i}{\partial d_j} \text{ and } \chi_{N,i\beta} = -\frac{\partial N_i}{\partial b_\beta}, \end{aligned} \tag{9}$$

where $v_{N,ij}$ and $\chi_{N,i\beta}$ are susceptibility matrices characterising the response of the abundance of consumer $i$ to perturbations in the death rate of consumer $j$ and outflux rate of resource $\beta$, respectively.

Repeating the same process with resource $\alpha$:

$$\begin{aligned} R_\alpha &\approx R_{\alpha\backslash 0} + \sum_{j=1}^{S} \underbrace{\frac{\partial R_\alpha}{\partial d_j}}_{v_{R,\alpha j}} \Delta d_j + \sum_{\beta=1}^{M} \underbrace{\frac{\partial R_\alpha}{\partial b_\beta}}_{-\chi_{R,\alpha\beta}} \Delta b_\beta \\ &\approx R_{\alpha\backslash 0} - \sum_{j=1}^{S} v_{R,\alpha j} \left[\frac{\sigma_g}{\sqrt{M}} z_{g,j0} R_0\right] - \sum_{\beta=1}^{M} \chi_{R,\alpha\beta} \left[\frac{\sigma_c}{\sqrt{M}} \left[\rho z_{g,0\beta} + \sqrt{1-\rho^2} z_{c,0\beta}\right] N_0\right], \\ &\text{where } v_{R,\alpha j} = \frac{\partial R_\alpha}{\partial d_j} \text{ and } \chi_{R,\alpha\beta} = -\frac{\partial R_\alpha}{\partial b_\beta}, \end{aligned} \tag{10}$$

where $v_{R,\alpha j}$ and $\chi_{R,\alpha\beta}$ are the susceptibility matrices characterising the response of the abundance of resource $\alpha$ to small perturbations in the death rate of consumer $j$ and supply rate of resource $\beta$, respectively.

Below, we collect all the four susceptibility matrices defined for the model for reference later, as:

$$v_{N,ij} = \frac{\partial N_i}{\partial d_j}, \quad v_{R,\alpha j} = \frac{\partial R_\alpha}{\partial d_j}, \quad \chi_{N,i\beta} = -\frac{\partial N_i}{\partial b_\beta}, \quad \chi_{R,\alpha\beta} = -\frac{\partial R_\alpha}{\partial b_\beta}. \tag{11}$$

#### A.3 Self-consistent dynamics of the cavity consumer and resource

The steady-state condition for the cavity consumer (with abundance $N_0$) and resource (with abundance $R_0$) are

$$\frac{\mathrm{d}N_0}{\mathrm{d}t} = 0 = N_0 \left[\mu_g \langle R\rangle + \frac{\sigma_g}{\sqrt{M}} \sum_{\alpha=1}^{M} z_{g,0\alpha} R_\alpha + \frac{\sigma_g}{\sqrt{M}} z_{g,00} R_0 - d_0\right] \tag{12}$$

and

$$\frac{\mathrm{d}R_0}{\mathrm{d}t} = 0 = I_0 - b_0 R_0 - R_0 \left[\mu_c \gamma^{-1} \langle N\rangle + \frac{\sigma_c}{\sqrt{M}} \sum_{i=1}^{S} \left[\rho z_{g,i0} + \sqrt{1-\rho^2} z_{c,i0}\right] N_i + \frac{\sigma_c}{\sqrt{M}} \left[\rho z_{g,00} + \sqrt{1-\rho^2} z_{c,00}\right] N_0\right]. \tag{13}$$

##### A.3.1 Cavity consumer equation

We first solve for the steady-state abundance of the cavity consumer $N_0$. Substituting expressions for $N_i$ and $R_\alpha$ from eq. (10), and plugging in the parameter sampling rules (Main text, Parametrisation) for $d_0$, we get:

$$\begin{aligned} 0 = N_0 \Bigg[ & \mu_g \langle R\rangle - \mu_d + \frac{\sigma_g}{\sqrt{M}} \sum_{\alpha=1}^{M} z_{g,0\alpha} R_{\alpha\backslash 0} - \frac{\sigma_g^2}{M} R_0 \sum_{\alpha=1}^{M} \sum_{j=1}^{S} v_{R,\alpha j} z_{g,0\alpha} z_{g,j0} \\ & - \frac{\sigma_g \sigma_c}{M} N_0 \sum_{\alpha=1}^{M} \sum_{\beta=1}^{M} \chi_{R,\alpha\beta} z_{g,0\alpha} \left[\rho z_{g,0\beta} + \sqrt{1-\rho^2} z_{c,0\beta}\right] + \frac{\sigma_g}{\sqrt{M}} z_{g,00} R_0 - \sigma_d z_{d,0} \Bigg]. \end{aligned} \tag{14}$$

To solve this equation and obtain the steady-state abundance $N_0$, we invoke the central limit theorem to express the many sums of weakly-correlated variables in eq. (14) as a single Gaussian random variable. The Gaussianity of the sums will appear due to the central limit theorem irrespective of whether the underlying $c_{i\alpha}$ follow uniform, Bernoulli or Gaussian distributions. To express these sums as a single Gaussian variable, we need to compute its first and second moments (which will contribute to the mean and fluctuations) in terms of the non-zero $M$-leading-order moments of the sums in eq. (14).

Most first moments in (14) are zero. The only non-zero term that contributes to the first moment is

$$\left\langle -\frac{\sigma_g \sigma_c}{M} N_0 \sum_{\alpha=1}^{M} \sum_{\beta=1}^{M} \chi_{R,\alpha\beta} z_{g,0\alpha} \left[\rho z_{g,0\beta} + \sqrt{1-\rho^2} z_{c,0\beta}\right] \right\rangle = -\sigma_g \sigma_c \rho \langle \chi_R \rangle N_0, \tag{15}$$

where $\langle \chi_R\rangle = 1/M \sum_{\alpha,\beta}^{M} \chi_{R,\alpha\beta} = 1/M \ \mathrm{Trace}(\chi_{R,\alpha\beta})$ in the large $M$ limit is the average diagonal entry of the susceptibility matrix $\chi_{R,\alpha\beta}$.

The non-zero, non-vanishing second moments are

$$\left\langle [-\sigma_d z_{0,d}]^2 \right\rangle = \sigma_d^2. \tag{16}$$

and

$$\left\langle \left[ \frac{\sigma_g}{\sqrt{M}} \sum_{\alpha=1}^{M} z_{g,0\alpha} R_{\alpha\backslash 0} \right]^2 \right\rangle = \sigma_g^2 \langle R^2 \rangle. \tag{17}$$

Note that here we have introduced the mean of the square of the steady-state resource abundance $\langle R^2 \rangle = \frac{1}{M} \sum_{\alpha=1}^{M} R_\alpha^2$, another unknown which we must later determine self-consistently.

One can show that the second moments of the other terms in (14), besides the final term, are sub-leading in $M$. (For a detailed discussion see [1] for GLV models and [2] for CRMs.

Combining the moments computed above, we can write a simplified version of Eq. (14):

$$0 = N_0 \left[ \mu_g \langle R \rangle - \mu_d - \sigma_g \sigma_c \rho \langle \chi_R \rangle N_0 + \sqrt{\sigma_g^2 \langle R^2 \rangle + \sigma_d^2} Z_N \right], \tag{18}$$

where $Z_N$ is a standard normal Gaussian variable. We can solve this equation to compute the cavity consumer's abundance ($N_0$) at steady state. There are two solutions that satisfy eq. (18)): When $N_0 = 0$ (extinction), or $N_0 = [\mu_g \langle R \rangle - \mu_d + \sqrt{\sigma_g^2 \langle R^2 \rangle + \sigma_d^2} Z_N] / \sigma_g \sigma_c \rho \langle \chi_R \rangle$ (survival). This means the cavity consumer abundance follows a normal distribution truncated at 0:

$$N_0 = \max \left\{ 0, \frac{\mu_g \langle R \rangle - \mu_d + \sqrt{\sigma_g^2 \langle R^2 \rangle + \sigma_d^2} Z_N}{\sigma_g \sigma_c \rho \langle \chi_R \rangle} \right\}. \tag{19}$$

We assign the probability a cavity consumer will survive ($N_0 > 0$) as $S^*/S = \phi_N$, which we will later determine self-consistently.

##### A.3.2 Cavity resource equation

We can now repeat similar steps to solve Eq. (13) to compute the steady-state abundance of resource 0. Substituting our linear response approximation for $N_i$ from Eq. (9) in eq. (13), we get:

$$\begin{aligned} 0 = \underbrace{\mu_I + \sigma_I z_{I,0}}_{I_0} - (\underbrace{\mu_b + \sigma_b z_{b,0}}_{b_0}) R_0 - R_0 \Bigg[ \mu_c \gamma^{-1} \langle N \rangle \\ + \frac{\sigma_c}{\sqrt{M}} \sum_{i=1}^{S} \left[ \rho z_{g,i0} + \sqrt{1-\rho^2} z_{c,i0} \right] \Bigg[ N_{i\backslash 0} - \sum_{j=1}^{S} v_{N,ij} \left[ \frac{\sigma_g}{\sqrt{M}} z_{g,j0} R_0 \right] \\ - \sum_{\beta=1}^{M} \chi_{N,i\beta} \left[ \frac{\sigma_c}{\sqrt{M}} \left[ \rho z_{g,0\beta} + \sqrt{1-\rho^2} z_{c,0\beta} \right] N_0 \right] \Bigg] \\ + \frac{\sigma_c}{\sqrt{M}} \left[ \rho z_{g,00} + \sqrt{1-\rho^2} z_{c,00} \right] N_0 \Bigg]. \end{aligned} \tag{20}$$

Similar to the cavity consumer, we need to compute the first and second non-vanishing moments of the terms in eq. (20) to express these sums as simplified random variables. Note that we cannot use the central limit theorem to combine the resource influx rate $I_0$ with consumption and dilution rates because it is mulitplied by $R_0$, so it is not considered in the following calculations.

The only term with a non-zero, leading order first moment is

$$\left\langle -R_0 \frac{\sigma_c}{\sqrt{M}} \sum_{i=1}^{S} \left[\rho z_{g,i0} + \sqrt{1-\rho^2} z_{c,i0}\right] \left[ -\sum_{j=1}^{S} v_{N,ij} \left[ \frac{\sigma_g}{\sqrt{M}} z_{g,j0} R_0 \right]\right] \right\rangle = \sigma_g \sigma_c \rho \gamma^{-1} \langle v_N \rangle R_0^2. \tag{21}$$

where $\langle v_N \rangle = 1/S \sum_{i,j}^{S} v_{N,ij} = 1/S \ \mathrm{Trace}(v_{N,ij})$ in the large $S$ limit is the average diagonal entry of the susceptibility matrix $v_{N,ij}$, and quantifies the average change in the abundance of consumer $i$ upon a small perturbation to its death rate.

The terms with non-zero, leading-order second moments are

$$\langle [-\sigma_b z_{b,0} R_0]^2 \rangle = \sigma_b^2 R_0^2 \tag{22}$$

and

$$\left\langle \left[ -R_0 \frac{\sigma_c}{\sqrt{M}} \sum_{i=1}^{S} \left[\rho z_{g,i0} + \sqrt{1-\rho^2} z_{c,i0}\right] N_{i\setminus 0} \right]^2 \right\rangle = \sigma_c^2 \gamma^{-1} \langle N^2 \rangle R_0^2. \tag{23}$$

Note that here we have introduced the mean of the square of the steady-state consumer abundance $\langle N^2 \rangle = \frac{1}{S}\sum_{i=1}^{S} N_i^2$, another unknown which we must later determine self-consistently.

Similar to the case of the cavity consumer, one can show that the second moments of the other sums and the cavity consumer fluctuation are sub-leading compared to the above terms.

We can now write the dynamics of the cavity resource in terms of the non-vanishing moments.

$$0 = [\mu_I + \sigma_I z_{I,0}] - \left[ \mu_b + \mu_c \gamma^{-1} \langle N \rangle + \sqrt{\sigma_c^2 \gamma^{-1} \langle N^2 \rangle + \sigma_b^2} Z_R \right] R_0 + \sigma_g \sigma_c \rho \gamma^{-1} \langle v_N \rangle R_0^2, \tag{24}$$

where the $-$ that would persist in $\sqrt{\sigma_c^2 \gamma^{-1} \langle N^2 \rangle + \sigma_b^2} Z_R$ has been absorbed into $Z_R$ for convenience.

Eq. (24) is a quadratic, meaning the solution for the cavity resource abundance $R_0$ is

$$\begin{aligned} R_0 =& \frac{\mu_b + \mu_c \gamma^{-1} \langle N \rangle + \sqrt{\sigma_c^2 \gamma^{-1} \langle N^2 \rangle + \sigma_b^2} Z_R}{2\sigma_c \sigma_g \rho \gamma^{-1} \langle v_N \rangle} \\ & \pm \frac{\sqrt{[\mu_b + \mu_c \gamma^{-1} \langle N \rangle + \sqrt{\sigma_c^2 \gamma^{-1} \langle N^2 \rangle + \sigma_b^2} Z_R]^2 - 4[\sigma_c \sigma_g \rho \gamma^{-1} \langle v_N \rangle][\mu_I + \sigma_I z_{I,0}]}}{2\sigma_c \sigma_g \rho \gamma^{-1} \langle v_N \rangle}. \end{aligned} \tag{25}$$

As we will show later on, $\langle v_N \rangle \leq 0$, so the only biologically relevant solution for $R_0$ is the negative solution.

Assuming that the means in the numerator of the second term in (25) are $\gg$ than their fluctuations, we can approximate the second term by taking its linear binomial expansion about its mean: This means $R_0$ can be approximated as

$$\begin{aligned} R_0 \approx \frac{1}{2\sigma_c \sigma_g \rho \gamma^{-1} \langle v_N \rangle} \Bigg[ & \mu_b + \mu_c \gamma^{-1} \langle N \rangle + \sqrt{\sigma_c^2 \gamma^{-1} \langle N^2 \rangle + \sigma_b^2} Z_R - \sqrt{[\mu_b + \mu_c \gamma^{-1} \langle N \rangle]^2 - 4\sigma_c \sigma_g \rho \gamma^{-1} \langle v_N \rangle \mu_I} \\ & - \frac{2[\mu_b + \mu_c \gamma^{-1} \langle N \rangle] \sqrt{\sigma_c^2 \gamma^{-1} \langle N^2 \rangle + \sigma_b^2} Z_R + [\sigma_c^2 \gamma^{-1} \langle N^2 \rangle + \sigma_b^2] Z_R^2 - 4\sigma_c \sigma_g \rho \gamma^{-1} \langle v_N \rangle \sigma_I z_{I,0}}{2\sqrt{[\mu_b + \mu_c \gamma^{-1} \langle N \rangle]^2 - 4\sigma_c \sigma_g \rho \gamma^{-1} \langle v_N \rangle \mu_I}} \Bigg]. \end{aligned} \tag{26}$$

#### A.4 Self-consistency equations

Since the cavity species and resource are statistically identical to the resident species and resources respectively, we invoke self-averaging to claim that the distribution of all consumer and resources in the steady-state community follow the same distributions as the cavity consumer and resource. Now, we derive the self-consistency equations describing the statistical moments of these abundance distributions: $\phi_N$, $\langle N\rangle$ and $\langle R\rangle$, $\langle N^2\rangle$ and $\langle R^2\rangle$, and $\langle v_N\rangle$ and $\langle\chi_R\rangle$.

##### A.4.1 The statistical moments of the consumer abundance distribution

Several of the unknowns are the moments of the consumer abundance distribution: the zeroth moment represents the consumer survival probability of consumer ($\phi_N$), the first moments represent the average consumer abundance $\langle N\rangle$, and similarly for the second moment $\langle N^2\rangle$ and $\langle R^2\rangle$. The $j^{\text{th}}$ moment of the zero-truncated Gaussian describing the consumer abundance distribution is

$$w_j\left(\mu_g\langle R\rangle-\mu_d,\sqrt{\sigma_g^2\langle R^2\rangle+\sigma_d^2}\right)=\left[\frac{\sqrt{\sigma_g^2\langle R^2\rangle+\sigma_d^2}}{\sigma_g\sigma_c\rho\langle\chi_R\rangle}\right]^j\int_{-\frac{\mu_g\langle R\rangle-\mu_d}{\sqrt{\sigma_g^2\langle R^2\rangle+\sigma_d^2}}}^{\infty}\frac{1}{\sqrt{2\pi}}\exp\left(-\frac{z^2}{2}\right)\left[z+\frac{\mu_g\langle R\rangle-\mu_d}{\sqrt{\sigma_g^2\langle R^2\rangle+\sigma_d^2}}\right]^j dz. \tag{27}$$

The 0th moment $\phi_N$ is

$$\phi_N=\int_{-\frac{\mu_g\langle R\rangle-\mu_d}{\sqrt{\sigma_g^2\langle R^2\rangle+\sigma_d^2}}}^{\infty}\frac{1}{\sqrt{2\pi}}\exp\left(-\frac{z^2}{2}\right)dz=\Phi\left(\frac{\mu_g\langle R\rangle-\mu_d}{\sqrt{\sigma_g^2\langle R^2\rangle+\sigma_d^2}}\right), \tag{28}$$

where $\Phi(...)$ is $1/2\times$the complementary error function.

The 1st moment $\langle N\rangle$ is

$$\langle N\rangle=\frac{\sqrt{\sigma_g^2\langle R^2\rangle+\sigma_d^2}}{\sigma_g\sigma_c\rho\langle\chi_R\rangle}\left[\frac{1}{\sqrt{2\pi}}\exp\left(-\frac{[\mu_g\langle R\rangle-\mu_d]^2}{2[\sigma_g^2\langle R^2\rangle+\sigma_d^2]}\right)+\frac{\mu_g\langle R\rangle-\mu_d}{\sqrt{\sigma_g^2\langle R^2\rangle+\sigma_d^2}}\,\Phi\left(\frac{\mu_g\langle R\rangle-\mu_d}{\sqrt{\sigma_g^2\langle R^2\rangle+\sigma_d^2}}\right)\right]. \tag{29}$$

The 2nd moment $\langle N^2\rangle$ is

$$\langle N^2\rangle=\left[\frac{\sqrt{\sigma_g^2\langle R^2\rangle+\sigma_d^2}}{\sigma_g\sigma_c\rho\langle\chi_R\rangle}\right]^2\left[\frac{\mu_g\langle R\rangle-\mu_d}{\sqrt{2\pi[\sigma_g^2\langle R^2\rangle+\sigma_d^2]}}\exp\left(-\frac{[\mu_g\langle R\rangle-\mu_d]^2}{2[\sigma_g^2\langle R^2\rangle+\sigma_d^2]}\right)\right.$$
$$\left.+\left[1+\frac{[\mu_g\langle R\rangle-\mu_d]^2}{\sigma_g^2\langle R^2\rangle+\sigma_d^2}\right]\Phi\left(\frac{\mu_g\langle R\rangle-\mu_d}{\sqrt{\sigma_g^2\langle R^2\rangle+\sigma_d^2}}\right)\right]. \tag{30}$$

##### A.4.2 The statistical moments of the resource abundance distribution

When resources are externally-supplied, their abundances do not follow a truncated Gaussian. To obtain the average resource abundance ($\langle R\rangle$) and its fluctuations ($\langle R^2\rangle$), we instead calculate the expected value of $R_0$ and $R_0^2$ using the expression for $R_0$ in eq. (26).

The 1st moment, or average resource abundance $\langle R\rangle$, is

$$\langle R\rangle = \frac{1}{2\sigma_c\sigma_g\rho\gamma^{-1}\langle v_N\rangle}\left[\mu_b + \mu_c\gamma^{-1}\langle N\rangle - \sqrt{[\mu_b + \mu_c\gamma^{-1}\langle N\rangle]^2 - 4\sigma_c\sigma_g\rho\gamma^{-1}\langle v_N\rangle\mu_I} - \frac{\sigma_c^2\gamma^{-1}\langle N^2\rangle + \sigma_b^2}{2\sqrt{[\mu_b + \mu_c\gamma^{-1}\langle N\rangle]^2 - 4\sigma_c\sigma_g\rho\gamma^{-1}\langle v_N\rangle\mu_I}}\right]. \tag{31}$$

The 2nd moment is

$$\begin{aligned}\langle R^2\rangle = \frac{1}{4[\sigma_g\sigma_c\rho\gamma^{-1}\langle v_N\rangle]^2}\Bigg[&2[\mu_b + \mu_c\gamma^{-1}\langle N\rangle]^2 + 2[\sigma_c^2\gamma^{-1}\langle N^2\rangle + \sigma_b^2] - 4\sigma_g\sigma_c\rho\gamma^{-1}\langle v_N\rangle\mu_I \\ &- 2[\mu_b + \mu_c\gamma^{-1}\langle N\rangle]\sqrt{[\mu_b + \mu_c\gamma^{-1}\langle N\rangle]^2 - 4\sigma_g\sigma_c\rho\gamma^{-1}\langle v_N\rangle\mu_I} \\ &- \frac{3[\mu_b + \mu_c\gamma^{-1}\langle N\rangle][\sigma_c^2\gamma^{-1}\langle N^2\rangle + \sigma_b^2]}{\sqrt{[\mu_b + \mu_c\gamma^{-1}\langle N\rangle]^2 - 4\sigma_g\sigma_c\rho\gamma^{-1}\langle v_N\rangle\mu_I}} \\ &+ \frac{4[\mu_b + \mu_c\gamma^{-1}\langle N\rangle]^2[\sigma_c^2\gamma^{-1}\langle N^2\rangle + \sigma_b^2] + 3[\sigma_c^2\gamma^{-1}\langle N^2\rangle + \sigma_b^2]^2 + 16[\sigma_g\sigma_c\rho\gamma^{-1}\langle v_N\rangle]^2\sigma_I^2}{4[[\mu_b + \mu_c\gamma^{-1}\langle N\rangle]^2 - 4\sigma_g\sigma_c\rho\gamma^{-1}\langle v_N\rangle\mu_I]}\Bigg].\end{aligned} \tag{32}$$

##### A.4.3 The susceptibilities

The susceptibility $\langle v_N\rangle$, which we remind ourselves equals $\langle\partial N_i/d_i\rangle$ or equivalently $\langle\partial N_0/\partial d_0\rangle$, is given by

$$\langle v_N\rangle = \left\langle \frac{\partial}{\partial d_0}\left(\max\left\{0, \frac{\mu_g\langle R\rangle - \mu_d + \sqrt{\sigma_g^2\langle R^2\rangle + \sigma_d^2}Z_N}{\sigma_g\sigma_c\rho\langle\chi_R\rangle}\right\}\right)\right\rangle \tag{33}$$

$$= -\frac{\phi_N}{\sigma_g\sigma_c\rho\langle\chi_R\rangle}. \tag{34}$$

The susceptibility $\langle\chi_R\rangle = -\langle\partial R_\alpha/\partial b_\alpha\rangle \equiv -\langle\partial R_0/\partial b_0\rangle$ is

$$\begin{aligned}\langle\chi_R\rangle = &-\left\langle\frac{\partial}{\partial b_0}\left(\frac{\mu_b + \mu_c\gamma^{-1}\langle N\rangle + \sqrt{\sigma_c^2\gamma^{-1}\langle N^2\rangle + \sigma_b^2}Z_R}{2\sigma_c\sigma_g\rho\gamma^{-1}\langle v_N\rangle} - \frac{\sqrt{[\mu_b + \mu_c\gamma^{-1}\langle N\rangle + \sqrt{\sigma_c^2\gamma^{-1}\langle N^2\rangle + \sigma_b^2}Z_R]^2 - 4[\sigma_c\sigma_g\rho\gamma^{-1}\langle v_N\rangle][\mu_I + \sigma_I z_{I,0}]}}{2\sigma_c\sigma_g\rho\gamma^{-1}\langle v_N\rangle}\right)\right\rangle. \\ \approx &-\frac{1}{2\sigma_c\sigma_g\rho\gamma^{-1}\langle v_N\rangle}\left[1 - \frac{\mu_b + \mu_c\gamma^{-1}\langle N\rangle}{\sqrt{[\mu_b + \mu_c\gamma^{-1}\langle N\rangle]^2 - 4\sigma_c\sigma_g\rho\gamma^{-1}\langle v_N\rangle\mu_I}} + \frac{3[\mu_b + \mu_c\gamma^{-1}\langle N\rangle][\sigma_c^2\gamma^{-1}\langle N^2\rangle + \sigma_b^2]}{2\left[[\mu_b + \mu_c\gamma^{-1}\langle N\rangle]^2 - 4\sigma_c\sigma_g\rho\gamma^{-1}\langle v_N\rangle\mu_I\right]^{\frac{3}{2}}}\right]\end{aligned} \tag{35}$$

#### A.5 Stability condition

Having derived properties of typical steady-state communities assembled from a pool of $S$ consumers and $M$ resources, we now proceed to derive when we expect these steady-states to be dynamically stable. For this, we will assess the sensitivity of the community steady-state to small perturbations in the abundances of all surviving consumers and resources at steady-state. (By perturbing only surviving consumers and all resources, we assume that our steady state is uninvadable.) When the distributions describing the sensitivities of consumers and resources to perturbations becomes ill-defined, the community becomes unstable. We will find that it is the second moment that becomes ill-defined, and the stability boundary thus corresponds to the condition for this second moment to diverge.

##### A.5.1 Sensitivity of consumer abundances to perturbations

From eq. (18), the steady-state abundance of the surviving cavity species $N_0^+$ can be written as

$$N_0^+ = \frac{1}{\sigma_g \sigma_c \rho \langle \chi_R \rangle} \left[ \mu_g \langle R \rangle - \mu_d + \frac{\sigma_g}{\sqrt{M}} \sum_{\alpha=1}^{M} z_{g,0\alpha} R_{\alpha \setminus 0} - \sigma_d z_{d,0} \right], \tag{36}$$

We now apply a random perturbation $\varepsilon \eta_\alpha$ to all resources $R_{\alpha\setminus 0}$: $R_{\alpha\setminus 0} \to R_{\alpha\setminus 0} + \varepsilon\eta_\alpha$ — where $\varepsilon \ll 1$ and $\eta_\alpha$ is a standard normal variable. After the perturbation, the steady-state abundance of the surviving cavity consumer is

$$N_0^+ = \frac{1}{\sigma_g \sigma_c \rho \langle \chi_R \rangle} \left[ \mu_g \langle R \rangle - \mu_d + \frac{\sigma_g}{\sqrt{M}} \sum_{\alpha=1}^{M} z_{g,0\alpha} \left[ R_{\alpha \setminus 0} + \varepsilon \eta_\alpha \right] - \sigma_d z_{d,0} \right]. \tag{37}$$

Next, we compute the sensitivity of this consumer $dN_0^+/d\varepsilon$ by taking its derivative with respect to the perturbation $\varepsilon$, given by:

$$\frac{\mathrm{d}N_0^+}{\mathrm{d}\varepsilon} = \frac{1}{\sigma_g \sigma_c \rho \langle \chi_R \rangle} \left[ \frac{\sigma_g}{\sqrt{M}} \sum_{\alpha=1}^{M} z_{g,0\alpha} \left[ \frac{\mathrm{d}R_{\alpha\setminus 0}}{\mathrm{d}\varepsilon} + \eta_\alpha \right] \right]. \tag{38}$$

The first moment $\langle dN_0^+/d\varepsilon \rangle = 0$. However, this is not the case for the second moment $\langle [dN_0^+/d\varepsilon]^2 \rangle$:

$$\left\langle \left[ \frac{\mathrm{d}N_0^+}{\mathrm{d}\varepsilon} \right]^2 \right\rangle = \frac{1}{[\sigma_g \sigma_c \rho \langle \chi_R \rangle]^2} \left\langle \left[ \frac{\sigma_g}{\sqrt{M}} \sum_{\alpha=1}^{M} z_{g,0\alpha} \left[ \frac{\mathrm{d}R_{\alpha\setminus 0}}{\mathrm{d}\varepsilon} + \eta_\alpha \right] \right]^2 \right\rangle \tag{39}$$

$$= \frac{1}{[\sigma_c \rho \langle \chi_R \rangle]^2} \left[ \left\langle \left[ \frac{\mathrm{d}R_{\alpha\setminus 0}}{d\varepsilon} \right]^2 \right\rangle + 1 \tag{40}$$

Because the cavity resource is statistically identical to all community members, we can substitute $\langle [R_{\alpha\setminus 0}/d\varepsilon]^2 \rangle$ with $\langle [R_0/d\varepsilon]^2 \rangle$:

$$\left\langle \left[ \frac{\mathrm{d}N_0^+}{d\varepsilon} \right]^2 \right\rangle = \frac{1}{[\sigma_c \rho \langle \chi_R \rangle]^2} \left[ \left\langle \left[ \frac{\mathrm{d}R_0}{d\varepsilon} \right]^2 \right\rangle + 1 \right]. \tag{41}$$

##### A.5.2 Sensitivity of resource abundances to perturbations

Starting with eq. (25), the cavity resource abundance $R_0$ can be described by the following equation:

$$R_0 = \frac{1}{2\sigma_c\sigma_g\rho\gamma^{-1}\langle v_N\rangle}\Bigg[\mu_b + \mu_c\gamma^{-1}\langle N\rangle + \frac{\sigma_c}{\sqrt{M}}\sum_{i=1}^{S^*}\left[\rho z_{g,i0} + \sqrt{1-\rho^2}z_{c,i0}\right]N^+_{i/0} + \sigma_b z_{b,0} - \Bigg[\Bigg[\mu_b + \mu_c\gamma^{-1}\langle N\rangle + \frac{\sigma_c}{\sqrt{M}}\sum_{i=1}^{S^*}\left[\rho z_{g,i0} + \sqrt{1-\rho^2}z_{c,i0}\right]N^+_{i/0} + \sigma_b z_{b,0}\Bigg]^2 - 4\sigma_c\sigma_g\rho\gamma^{-1}\langle v_N\rangle[\mu_I + \sigma_I z_{I,0}]\Bigg]^{\frac{1}{2}}\Bigg]. \tag{42}$$

We now perturb surviving consumers abundances in a similar fashion to surviving resources in eq. (37) – $N^+_{i\backslash 0} \to N^+_{i\backslash 0} + \varepsilon\eta_i$ — giving us

$$R_0 = \frac{1}{2\sigma_c\sigma_g\rho\gamma^{-1}\langle v_N\rangle}\Bigg[\mu_b + \mu_c\gamma^{-1}\langle N\rangle + \frac{\sigma_c}{\sqrt{M}}\sum_{i=1}^{S^*}\left[\rho z_{g,i0} + \sqrt{1-\rho^2}z_{c,i0}\right]\left[N^+_{i/0} + \varepsilon\eta_i\right] + \sigma_b z_{b,0} - \Bigg[\Bigg[\mu_b + \mu_c\gamma^{-1}\langle N\rangle + \frac{\sigma_c}{\sqrt{M}}\sum_{i=1}^{S^*}\left[\rho z_{g,i0} + \sqrt{1-\rho^2}z_{c,i0}\right]\left[N^+_{i/0} + \varepsilon\eta_i\right] + \sigma_b z_{b,0}\Bigg]^2 - 4\sigma_c\sigma_g\rho\gamma^{-1}\langle v_N\rangle[\mu_I + \sigma_I z_{I,0}]\Bigg]^{\frac{1}{2}}\Bigg]. \tag{43}$$

The cavity resource's sensitivity to perturbations is

$$\frac{\mathrm{d}R_0}{\mathrm{d}\varepsilon} = \frac{1}{2\sigma_c\sigma_g\rho\gamma^{-1}\langle v_N\rangle}\Bigg[\frac{\sigma_c}{\sqrt{M}}\sum_{i=1}^{S^*}\left[\rho z_{g,i0} + \sqrt{1-\rho^2}z_{c,i0}\right]\left[\frac{\mathrm{d}N^+_{i/0}}{\mathrm{d}\varepsilon} + \eta_i\right] - \frac{\mathrm{d}}{\mathrm{d}\varepsilon}\Bigg[\Bigg[\mu_b + \mu_c\gamma^{-1}\langle N\rangle + \frac{\sigma_c}{\sqrt{M}}\sum_{i=1}^{S^*}\left[\rho z_{g,i0} + \sqrt{1-\rho^2}z_{c,i0}\right]\left[N^+_{i/0} + \varepsilon\eta_i\right] + \sigma_b z_{b,0}\Bigg]^2 - 4\sigma_c\sigma_g\rho\gamma^{-1}\langle v_N\rangle[\mu_I + \sigma_I z_{I,0}]\Bigg]^{\frac{1}{2}}\Bigg]. \tag{44}$$

To differentiate the last term of eq. (44), we denote $f(\varepsilon) = \mu_b + \mu_c\gamma^{-1}\langle N\rangle + [\sigma_c/\sqrt{M}]\sum_{i=1}^{S^*}[\rho z_{g,i0} + \sqrt{1-\rho^2}z_{c,i0}][N^+_{i/0} + \varepsilon\eta_i] + \sigma_b z_{b,0}$. Therefore, the final term can be reduced to

$$\begin{aligned} -\frac{\mathrm{d}}{\mathrm{d}\varepsilon}[f(\varepsilon)^2 - 4\sigma_c\sigma_g\rho\gamma^{-1}\langle v_N\rangle[\mu_I + \sigma_I z_{I,0}]]^{\frac{1}{2}} &= -\frac{1}{2[f(\varepsilon)^2 - 4\sigma_c\sigma_g\rho\gamma^{-1}\langle v_N\rangle[\mu_I + \sigma_I z_{I,0}]]^{\frac{1}{2}}} \times 2f(\varepsilon)\frac{\mathrm{d}f(\varepsilon)}{\mathrm{d}\varepsilon} \\ &= -\frac{f(\varepsilon)\sigma_c\sum_{i=1}^{S^*}\left[\rho z_{g,i0} + \sqrt{1-\rho^2}z_{c,i0}\right]\left[\frac{\mathrm{d}N^+_{i/0}}{\mathrm{d}\varepsilon} + \eta_i\right]}{\sqrt{M}[f(\varepsilon)^2 - 4\sigma_c\sigma_g\rho\gamma^{-1}\langle v_N\rangle[\mu_I + \sigma_I z_{I,0}]]^{\frac{1}{2}}} \end{aligned} \tag{45}$$

Therefore, $\mathrm{d}R_0/\mathrm{d}\varepsilon$ is

$$\frac{\mathrm{d}R_0}{\mathrm{d}\varepsilon} = \frac{1}{2\sigma_c\sigma_g\rho\gamma^{-1}\langle v_N\rangle}\left[\frac{\sigma_c}{\sqrt{M}}\sum_{i=1}^{S^*}\left[\rho z_{g,i0} + \sqrt{1-\rho^2}z_{c,i0}\right]\left[\frac{\mathrm{d}N^+_{i/0}}{\mathrm{d}\varepsilon} + \eta_i\right]\right]\left[1 - \frac{f(\varepsilon)}{[f(\varepsilon)^2 - 4\sigma_c\sigma_g\rho\gamma^{-1}\langle v_N\rangle[\mu_I + \sigma_I z_{I,0}]]^{\frac{1}{2}}}\right]. \tag{46}$$

The second moment of the sensitivity to perturbations, $\langle [dR_0^+/d\varepsilon]^2\rangle$, is

$$
\begin{aligned}
\left\langle \left[\frac{\mathrm{d}R_0}{\mathrm{d}\varepsilon}\right]^2 \right\rangle =& \frac{1}{4[\sigma_c\sigma_g\rho\gamma^{-1}\langle v_N\rangle]^2} \left\langle \left[\frac{\sigma_c}{\sqrt{M}}\sum_{i=1}^{S^*}\left[\rho z_{g,i0}+\sqrt{1-\rho^2}z_{c,i0}\right]\left[\frac{\mathrm{d}N_{i/0}^+}{\mathrm{d}\varepsilon}+\eta_i\right]\right]^2 \right. \\
& \left[1-\frac{2f(\varepsilon)}{[f(\varepsilon)^2-4\sigma_c\sigma_g\rho\gamma^{-1}\langle v_N\rangle[\mu_I+\sigma_I z_{I,0}]]^{\frac{1}{2}}} \right. \\
& \left.\left. +\frac{f(\varepsilon)^2}{f(\varepsilon)^2-4\sigma_c\sigma_g\rho\gamma^{-1}\langle v_N\rangle[\mu_I+\sigma_I z_{I,0}]}\right]\right\rangle \\
\approx& \frac{1}{4\left[\sigma_c\sigma_g\rho\gamma^{-1}\langle v_N\rangle\right]^2} \left\langle \left[\frac{\sigma_c}{\sqrt{M}}\sum_{i=1}^{S^*}\left[\rho z_{g,i0}+\sqrt{1-\rho^2}z_{c,i0}\right]\left[\frac{\mathrm{d}N_{i/0}^+}{\mathrm{d}\varepsilon}+\eta_i\right]\right]^2 \right\rangle \\
& \left\langle \left[1-\frac{2f(\varepsilon)}{[f(\varepsilon)^2-4\sigma_c\sigma_g\rho\gamma^{-1}\langle v_N\rangle[\mu_I+\sigma_I z_{I,0}]]^{\frac{1}{2}}} \right.\right. \\
& \left.\left. +\frac{f(\varepsilon)^2}{f(\varepsilon)^2-4\sigma_c\sigma_g\rho\gamma^{-1}\langle v_N\rangle[\mu_I+\sigma_I z_{I,0}]}\right]\right\rangle .
\end{aligned}
\tag{47}
$$

We can treat the two terms in the angle brackets as approximately independent because their correlation is subleading in the large-$M$ limit. The expectation of the first term is

$$
\left\langle \left[\frac{\sigma_c}{\sqrt{M}}\sum_{i=1}^{S^*}\left[\rho z_{g,i0}+\sqrt{1-\rho^2}z_{c,i0}\right]\left[\frac{\mathrm{d}N_{i/0}^+}{\mathrm{d}\varepsilon}+\eta_i\right]\right]^2 \right\rangle = \sigma_c^2\phi_N\gamma^{-1}\left[\left\langle\left[\frac{\mathrm{d}N_{i/0}^+}{\mathrm{d}\varepsilon}\right]^2\right\rangle+1\right]. \tag{48}
$$

Turning to the second term in eq. (47), we cannot compute it in its current form, as it is a non-linear function of $f(\varepsilon)$. To linearise the expression in terms of $f(\varepsilon)$, we note that $f(\varepsilon)^2-4\sigma_c\sigma_g\rho\gamma^{-1}\langle v_N\rangle[\mu_I+\sigma_I z_{I,0}]$ is composed of means and fluctuations:

$$
\begin{aligned}
&\left[\mu_b+\mu_c\gamma^{-1}\langle N\rangle+\frac{\sigma_c}{\sqrt{M}}\sum_{i=1}^{S^*}\left[\rho z_{g,i0}+\sqrt{1-\rho^2}z_{c,i0}\right]\left[N_{i/0}^+ +\varepsilon\eta_i\right]+\sigma_b z_{b,0}\right]^2-4\sigma_c\sigma_g\rho\gamma^{-1}\langle v_N\rangle[\mu_I+\sigma_I z_{I,0}] \\
&=\underbrace{\left[\mu_b+\mu_c\gamma^{-1}\langle N\rangle\right]^2-4\sigma_c\sigma_g\rho\gamma^{-1}\langle v_N\rangle\mu_I}_{\text{means}} \\
&\quad \underbrace{\begin{aligned}&+2\left[\mu_b+\mu_c\gamma^{-1}\langle N\rangle\right]\left[\frac{\sigma_c}{\sqrt{M}}\sum_{i=1}^{S^*}\left[\rho z_{g,i0}+\sqrt{1-\rho^2}z_{c,i0}\right]\left[N_{i/0}^+ +\varepsilon\eta_i\right]+\sigma_b z_{b,0}\right] \\ &+\left[\frac{\sigma_c}{\sqrt{M}}\sum_{i=1}^{S^*}\left[\rho z_{g,i0}+\sqrt{1-\rho^2}z_{c,i0}\right]\left[N_{i/0}^+ +\varepsilon\eta_i\right]+\sigma_b z_{b,0}\right]^2-4\sigma_c\sigma_g\rho\gamma^{-1}\langle v_N\rangle\sigma_I z_{I,0}\,.\end{aligned}}_{\text{fluctuations}}
\end{aligned}
\tag{49}
$$

For shorthand, we will rewrite these as

$$
\underbrace{\mu(\varepsilon)^2-4F\mu_I}_{\text{means}}+\underbrace{2\mu\sigma(\varepsilon)+\sigma(\varepsilon)^2-4F\sigma_I z_{I,0}}_{\text{fluctuations}}\,. \tag{50}
$$

where $\mu(\varepsilon)=\mu_b+\mu_c\gamma^{-1}\langle N\rangle$, $F=\sigma_c\sigma_g\rho\gamma^{-1}\langle v_N\rangle$ and $\sigma(\varepsilon)=\frac{\sigma_c}{\sqrt{M}}\sum_{i=1}^{S^*}\left[\rho z_{g,i0}+\sqrt{1-\rho^2}z_{c,i0}\right]\left[N_{i/0}^+ +\varepsilon\eta_i\right]+\sigma_b z_{b,0}$.

Assuming that the means are far larger than the fluctuations, we can approximate this term by taking the linear binomial expansion about its mean. Therefore,

$$\begin{aligned}
-\frac{2f(\varepsilon)}{[f(\varepsilon)^2-4\sigma_c\sigma_g\rho\gamma^{-1}\langle v_N\rangle[\mu_I+\sigma_I z_{I,0}]]^{\frac{1}{2}}} &\approx -\frac{2f(\varepsilon)}{[\mu(\varepsilon)^2-4F\mu_I]^{\frac{1}{2}}}\left[1-\frac{2\mu\sigma(\varepsilon)+\sigma(\varepsilon)^2-4F\sigma_I z_{I,0}}{2\left[\mu(\varepsilon)^2-4F\mu_I\right]}\right] \\
&\approx -\frac{2\left[\mu+\sigma(\varepsilon)\right]}{[\mu(\varepsilon)^2-4F\mu_I]^{\frac{1}{2}}}\left[1-\frac{2\mu\sigma(\varepsilon)+\sigma(\varepsilon)^2-4F\sigma_I z_{I,0}}{2\left[\mu(\varepsilon)^2-4F\mu_I\right]}\right]
\end{aligned} \tag{51}$$

and

$$\begin{aligned}
\frac{f(\varepsilon)^2}{f(\varepsilon)^2-4\sigma_c\sigma_g\rho\gamma^{-1}\langle v_N\rangle[\mu_I+\sigma_I z_{I,0}]} &\approx \frac{f(\varepsilon)^2}{\mu(\varepsilon)^2-4F\mu_I}\left[1-\frac{2\mu\sigma(\varepsilon)+\sigma(\varepsilon)^2-4F\sigma_I z_{I,0}}{\mu(\varepsilon)^2-4F\mu_I}\right] \\
&\approx \frac{[\mu+\sigma(\varepsilon)]^2}{\mu(\varepsilon)^2-4F\mu_I}\left[1-\frac{2\mu\sigma(\varepsilon)+\sigma(\varepsilon)^2-4F\sigma_I z_{I,0}}{\mu(\varepsilon)^2-4F\mu_I}\right].
\end{aligned} \tag{52}$$

Hence, the second term in angle brackets in eq. (47) is approximately

$$\begin{aligned}
&\left\langle 1-\frac{2\left[\mu+\sigma(\varepsilon)\right]}{[\mu(\varepsilon)^2-4F\mu_I]^{\frac{1}{2}}}\left[1-\frac{2\mu\sigma(\varepsilon)+\sigma(\varepsilon)^2-4F\sigma_I z_{I,0}}{2\left[\mu(\varepsilon)^2-4F\mu_I\right]}\right]+\frac{[\mu+\sigma(\varepsilon)]^2}{\mu(\varepsilon)^2-4F\mu_I}\left[1-\frac{2\mu\sigma(\varepsilon)+\sigma(\varepsilon)^2-4F\sigma_I z_{I,0}}{\mu(\varepsilon)^2-4F\mu_I}\right]\right\rangle. \\
&\approx 1-\frac{2}{[\mu(\varepsilon)^2-4F\mu_I]^{\frac{1}{2}}}\left[\mu-\frac{\mu\left[2\mu\langle\sigma(\varepsilon)\rangle+\langle\sigma(\varepsilon)^2\rangle-4F\sigma_I\langle z_{I,0}\rangle\right]}{2\left[\mu(\varepsilon)^2-4F\mu_I\right]}+\langle\sigma(\varepsilon)\rangle-\frac{2\mu\langle\sigma(\varepsilon)^2\rangle+\langle\sigma(\varepsilon)^3\rangle-4F\sigma_I\langle\sigma(\varepsilon)\rangle\langle z_{I,0}\rangle}{2\left[\mu(\varepsilon)^2-4F\mu_I\right]}\right] \\
&+\frac{1}{\mu(\varepsilon)^2-4F\mu_I}\Bigg[\mu^2-\frac{\mu^2\left[2\mu\langle\sigma(\varepsilon)\rangle+\langle\sigma(\varepsilon)^2\rangle-4F\sigma_I\langle z_{I,0}\rangle\right]}{\mu(\varepsilon)^2-4F\mu_I}+2\mu\langle\sigma(\varepsilon)\rangle \\
&\qquad -\frac{2\mu\left[2\mu\langle\sigma(\varepsilon)^2\rangle+\langle\sigma(\varepsilon)^3\rangle-4F\sigma_I\langle\sigma(\varepsilon)\rangle\langle z_{I,0}\rangle\right]}{\mu(\varepsilon)^2-4F\mu_I}+\langle\sigma(\varepsilon)^2\rangle-\frac{2\mu\langle\sigma(\varepsilon)^3\rangle+\langle\sigma(\varepsilon)^4\rangle-4F\sigma_I\langle\sigma(\varepsilon)^2\rangle\langle z_{I,0}\rangle}{\mu(\varepsilon)^2-4F\mu_I}\Bigg].
\end{aligned} \tag{53}$$

This expression has terms including the 1st to the 4th moment of $\sigma(\varepsilon)$. For ease, we will calculate the non-zero components of these moments below, and later plug them into the terms in eq. (53):

$$\langle\sigma(\varepsilon)\rangle=\frac{\sigma_c}{\sqrt{M}}\sum_{i=1}^{S^*}\left[\rho\langle z_{g,i0}\rangle+\sqrt{1-\rho^2}\langle z_{c,i0}\rangle\right]\left[\langle N_{i/0}^+\rangle+\varepsilon\langle\eta_i\rangle\right]+\sigma_b\langle z_{b,0}\rangle=0. \tag{54}$$

$$\begin{aligned}
\langle\sigma(\varepsilon)^2\rangle &= \frac{\sigma_c^2}{M}\sum_{i}^{S^*}\left[\rho^2\langle z_{g,i0}^2\rangle+2\rho\sqrt{1-\rho^2}\langle z_{g,i0}z_{c,i0}\rangle+[1-\rho^2]\langle z_{c,i0}^2\rangle\right]\left[\langle {N_{i/0}^+}^2\rangle+2\varepsilon\langle N_{i/0}^+\rangle\langle\eta_i\rangle+\varepsilon^2\langle\eta_i^2\rangle\right] \\
&\quad+\sigma_b^2\langle z_{b,0}^2\rangle \\
&= \sigma_c^2\gamma^{-1}\phi_N\left[\frac{\langle N^2\rangle}{\phi_N}+\varepsilon^2\right]+\sigma_b^2 \\
&\approx \sigma_c^2\gamma^{-1}\langle N^2\rangle+\sigma_b^2
\end{aligned} \tag{55}$$

since $0<\varepsilon\ll 1$.

$$\langle\sigma(\varepsilon)^3\rangle=0. \tag{56}$$

$$
\begin{aligned}
\langle\sigma(\varepsilon)^4\rangle =&\frac{\sigma_c^4}{M^2}\left[\sum_i^{S^*}\left[\rho^4\langle z_{g,i0}^4\rangle + 4\rho^3\sqrt{1-\rho^2}\langle z_{g,i0}^3\rangle\langle z_{c,i0}\rangle + 6\rho^2[1-\rho^2]\langle z_{g,i0}^2\rangle\langle z_{c,i0}^2\rangle\right.\right.\\
&\qquad\left. + 4\rho[1-\rho^2]^{\frac{3}{2}}\langle z_{g,i0}\rangle\langle z_{c,i0}^3\rangle + [1-\rho^2]^2\langle z_{c,i0}^4\rangle\right]\\
&\qquad\left[\langle {N_{i/0}^+}^4\rangle + 4\varepsilon\langle {N_{i/0}^+}^3\rangle\langle\eta_i\rangle + 6\varepsilon^2\langle {N_{i/0}^+}^2\rangle\langle\eta_i^2\rangle + 4\varepsilon^3\langle N_{i/0}^+\rangle\langle\eta_i^3\rangle + \varepsilon^4\langle\eta_i^4\rangle\right]\\
&\qquad + 3\sum_{i,j\neq i}^{S^*}\left[\rho^2\langle z_{g,i0}^2\rangle + 2\rho\sqrt{1-\rho^2}\langle z_{g,i0}z_{c,i0}\rangle + [1-\rho^2]\langle z_{c,i0}^2\rangle\right]\\
&\qquad\quad\left[\rho^2\langle z_{g,j0}^2\rangle + 2\rho\sqrt{1-\rho^2}\langle z_{g,j0}z_{c,j0}\rangle + [1-\rho^2]\langle z_{c,j0}^2\rangle\right]\\
&\qquad\quad\left.\left[\langle {N_{i/0}^+}^2\rangle + 2\varepsilon\langle N_{i/0}^+\rangle\langle\eta_i\rangle + \varepsilon^2\langle\eta_i^2\rangle\right]\left[\langle {N_{j/0}^+}^2\rangle + 2\varepsilon\langle N_{j/0}^+\rangle\langle\eta_j\rangle + \varepsilon^2\langle\eta_j^2\rangle\right]\right]\\
&+\frac{6\sigma_c^2\sigma_b^2}{M}\sum_i^{S^*}\left[\rho^2\langle z_{g,i0}^2\rangle + 2\rho\sqrt{1-\rho^2}\langle z_{g,i0}z_{c,i0}\rangle + [1-\rho^2]\langle z_{c,i0}^2\rangle\right]\\
&\qquad\quad\left[\langle {N_{i/0}^+}^2\rangle + 2\varepsilon\langle N_{i/0}^+\rangle\langle\eta_i\rangle + \varepsilon^2\langle\eta_i^2\rangle\right]\langle z_{I,0}^2\rangle\\
&+\sigma_b^4\langle z_{I,0}^4\rangle\\
=&\frac{\sigma_c^4\phi_N S}{M^2}\left[3\rho^4+6\rho^2-6\rho^4+3-6\rho^2+3\rho^4\right]\left[\langle {N_{i/0}^+}^4\rangle + 6\varepsilon^2\frac{\langle N^2\rangle}{\phi_N}+3\varepsilon^4\right]\\
&+\frac{\sigma_c^4\phi_N^2S^2}{M^2}\left[\rho^2+1-\rho^2\right]^2\left[\frac{\langle N^2\rangle}{\phi_N}+\varepsilon^2\right]^2+\frac{6\sigma_c^2\sigma_b^2\phi_N S}{M}\left[\rho^2+1-\rho^2\right]\left[\frac{\langle N^2\rangle}{\phi_N}+\varepsilon^2\right]+3\sigma_b^4\\
\approx&3\left[\sigma_c^2\phi_N\gamma^{-1}\right]^2\left[\frac{\langle N^2\rangle}{\phi_N}\right]^2+6\sigma_c^2\sigma_b^2\phi_N\gamma^{-1}\left[\frac{\langle N^2\rangle}{\phi_N}\right]+3\sigma_b^4\\
\approx&3\left[\sigma_c^2\gamma^{-1}\langle N^2\rangle\right]^2+6\sigma_b^2\left[\sigma_c^2\gamma^{-1}\langle N^2\rangle\right]+3\sigma_b^4\\
\approx&3\left[\sigma_c^2\gamma^{-1}\langle N^2\rangle+\sigma_b^2\right]^2.
\end{aligned}
\tag{57}
$$

as $0 < \varepsilon \ll 1$ and $M \to \infty$.

Returning to eq. (53) plugging in our moments of $\sigma(\varepsilon)$, as well as $\mu(\varepsilon)$ and $F$ reduces the expression to

$$
\begin{aligned}
&1-\frac{2}{[\mu(\varepsilon)^2-4F\mu_I]^{\frac{1}{2}}}\left[\mu-\frac{3\mu\langle\sigma(\varepsilon)^2\rangle}{2\left[\mu(\varepsilon)^2-4F\mu_I\right]}\right]+\frac{1}{\mu(\varepsilon)^2-4F\mu_I}\left[\mu^2+\langle\sigma(\varepsilon)^2\rangle-\frac{5\mu^2\langle\sigma(\varepsilon)^2\rangle+\langle\sigma(\varepsilon)^4\rangle}{\mu(\varepsilon)^2-4F\mu_I}\right]\\
&\approx 1-\frac{2}{\left[\left[\mu_b+\mu_c\gamma^{-1}\langle N\rangle\right]^2-4\sigma_c\sigma_g\rho\gamma^{-1}\langle v_N\rangle\mu_I\right]^{\frac{1}{2}}}\left[\mu_b+\mu_c\gamma^{-1}\langle N\rangle-\frac{3\left[\mu_b+\mu_c\gamma^{-1}\langle N\rangle\right]\left[\sigma_c^2\gamma^{-1}\langle N^2\rangle+\sigma_b^2\right]}{2\left[\left[\mu_b+\mu_c\gamma^{-1}\langle N\rangle\right]^2-4\sigma_c\sigma_g\rho\gamma^{-1}\langle v_N\rangle\mu_I\right]}\right]\\
&\quad+\frac{1}{\left[\mu_b+\mu_c\gamma^{-1}\langle N\rangle\right]^2-4\sigma_c\sigma_g\rho\gamma^{-1}\langle v_N\rangle\mu_I}\left[\left[\mu_b+\mu_c\gamma^{-1}\langle N\rangle\right]^2+\sigma_c^2\gamma^{-1}\langle N^2\rangle+\sigma_b^2\right.\\
&\qquad-\frac{5\left[\mu_b+\mu_c\gamma^{-1}\langle N\rangle\right]^2\left[\sigma_c^2\gamma^{-1}\langle N^2\rangle+\sigma_b^2\right]}{\left[\mu_b+\mu_c\gamma^{-1}\langle N\rangle\right]^2-4\sigma_c\sigma_g\rho\gamma^{-1}\langle v_N\rangle\mu_I}\\
&\qquad\left.-\frac{3\left[\sigma_c^2\gamma^{-1}\langle N^2\rangle+\sigma_b^2\right]^2}{\left[\mu_b+\mu_c\gamma^{-1}\langle N\rangle\right]^2-4\sigma_c\sigma_g\rho\gamma^{-1}\langle v_N\rangle\mu_I}\right].
\end{aligned}
\tag{58}
$$

Therefore, the final approximation for $\langle[\mathrm{d}R_0/\mathrm{d}\varepsilon]^2\rangle$ is

$$\left\langle\left[\frac{\mathrm{d}R_0}{\mathrm{d}\varepsilon}\right]^2\right\rangle \approx \frac{\sigma_c^2\phi_N\gamma^{-1}}{4\left[\sigma_c\sigma_g\rho\gamma^{-1}\langle v_N\rangle\right]^2}\left[\left\langle\left[\frac{\mathrm{d}N_{i/0}^+}{\mathrm{d}\varepsilon}\right]^2\right\rangle+1\right]$$
$$\times\left[1-\frac{2}{\left[\left[\mu_b+\mu_c\gamma^{-1}\langle N\rangle\right]^2-4\sigma_c\sigma_g\rho\gamma^{-1}\langle v_N\rangle\mu_I\right]^{\frac{1}{2}}}\left[\mu_b+\mu_c\gamma^{-1}\langle N\rangle-\frac{3\left[\mu_b+\mu_c\gamma^{-1}\langle N\rangle\right]\left[\sigma_c^2\gamma^{-1}\langle N^2\rangle+\sigma_b^2\right]}{2\left[\left[\mu_b+\mu_c\gamma^{-1}\langle N\rangle\right]^2-4\sigma_c\sigma_g\rho\gamma^{-1}\langle v_N\rangle\mu_I\right]}\right]\right.$$
$$+\frac{1}{\left[\mu_b+\mu_c\gamma^{-1}\langle N\rangle\right]^2-4\sigma_c\sigma_g\rho\gamma^{-1}\langle v_N\rangle\mu_I}\left[\left[\mu_b+\mu_c\gamma^{-1}\langle N\rangle\right]^2+\sigma_c^2\gamma^{-1}\langle N^2\rangle+\sigma_b^2-\frac{5\left[\mu_b+\mu_c\gamma^{-1}\langle N\rangle\right]^2\left[\sigma_c^2\gamma^{-1}\langle N^2\rangle+\sigma_b^2\right]}{\left[\mu_b+\mu_c\gamma^{-1}\langle N\rangle\right]^2-4\sigma_c\sigma_g\rho\gamma^{-1}\langle v_N\rangle\mu_I}\right.$$
$$\left.\left.-\frac{3\left[\sigma_c^2\gamma^{-1}\langle N^2\rangle+\sigma_b^2\right]^2}{\left[\mu_b+\mu_c\gamma^{-1}\langle N\rangle\right]^2-4\sigma_c\sigma_g\rho\gamma^{-1}\langle v_N\rangle\mu_I}\right]\right]. \tag{59}$$

Because the cavity consumer is statistically identical to all community members, we can substitute $\langle[\mathrm{d}N_{i\backslash 0}^+/\mathrm{d}\varepsilon]^2\rangle$ with $\langle[\mathrm{d}N_0/\mathrm{d}\varepsilon]^2\rangle$:

$$\left\langle\left[\frac{\mathrm{d}R_0}{\mathrm{d}\varepsilon}\right]^2\right\rangle \approx \frac{\phi_N\gamma^{-1}}{4\left[\sigma_g\rho\gamma^{-1}\langle v_N\rangle\right]^2}\left[\left\langle\left[\frac{\mathrm{d}N_0^+}{\mathrm{d}\varepsilon}\right]^2\right\rangle+1\right]$$
$$\times\left[1-\frac{2}{\left[\left[\mu_b+\mu_c\gamma^{-1}\langle N\rangle\right]^2-4\sigma_c\sigma_g\rho\gamma^{-1}\langle v_N\rangle\mu_I\right]^{\frac{1}{2}}}\left[\mu_b+\mu_c\gamma^{-1}\langle N\rangle-\frac{3\left[\mu_b+\mu_c\gamma^{-1}\langle N\rangle\right]\left[\sigma_c^2\gamma^{-1}\langle N^2\rangle+\sigma_b^2\right]}{2\left[\left[\mu_b+\mu_c\gamma^{-1}\langle N\rangle\right]^2-4\sigma_c\sigma_g\rho\gamma^{-1}\langle v_N\rangle\mu_I\right]}\right]\right.$$
$$+\frac{1}{\left[\mu_b+\mu_c\gamma^{-1}\langle N\rangle\right]^2-4\sigma_c\sigma_g\rho\gamma^{-1}\langle v_N\rangle\mu_I}\left[\left[\mu_b+\mu_c\gamma^{-1}\langle N\rangle\right]^2+\sigma_c^2\gamma^{-1}\langle N^2\rangle+\sigma_b^2-\frac{5\left[\mu_b+\mu_c\gamma^{-1}\langle N\rangle\right]^2\left[\sigma_c^2\gamma^{-1}\langle N^2\rangle+\sigma_b^2\right]}{\left[\mu_b+\mu_c\gamma^{-1}\langle N\rangle\right]^2-4\sigma_c\sigma_g\rho\gamma^{-1}\langle v_N\rangle\mu_I}\right.$$
$$\left.\left.-\frac{3\left[\sigma_c^2\gamma^{-1}\langle N^2\rangle+\sigma_b^2\right]^2}{\left[\mu_b+\mu_c\gamma^{-1}\langle N\rangle\right]^2-4\sigma_c\sigma_g\rho\gamma^{-1}\langle v_N\rangle\mu_I}\right]\right]. \tag{60}$$

##### A.5.3 Stability condition

Solving for $\langle\left[\mathrm{d}N_0^+/d\varepsilon\right]^2\rangle$ and $\langle[\mathrm{d}R_0/\mathrm{d}\varepsilon]^2\rangle$ gives us expressions of the form $\langle[\mathrm{d}N_0^+/d\varepsilon]^2\rangle = [B+1]/[AB-1]$ and $\langle[\mathrm{d}R_0/\mathrm{d}\varepsilon]^2\rangle = [A+1]/[AB-1]$, where $1/A = 1/[\sigma_c\rho\langle\chi_R\rangle]^2$ and $1/B = [\phi_N\gamma^{-1}/4[\sigma_g\gamma^{-1}\rho\langle v_N\rangle]^2] \times$ [the long expression in $\langle[\mathrm{d}R_0/\mathrm{d}\varepsilon]^2\rangle$] Communities undergo a transition to instability when these sensitivities diverge, which occurs when $AB-1=0$. Therefore, communities are stable when $AB-1>0$ or equivalently when $1/AB<1$:

$$\frac{\phi_N\gamma^{-1}}{4[\sigma_c\rho\langle\chi_R\rangle]^2[\sigma_g\gamma^{-1}\rho\langle v_N\rangle]^2}\left[\ldots\right]<1. \tag{61}$$

We can simplify the expression outside the bracket by plugging in the expression for $\langle v_N \rangle$ in eq. (34), which reduces the expression to

$$\frac{1}{4\rho^2\phi_N\gamma^{-1}}\left[...\right] < 1, \text{ or equivalently, } \frac{1}{4\rho^2}\left[...\right] < \phi_N\gamma^{-1}. \tag{62}$$

Therefore, the full stability condition is

**Stability condition for the externally-supplied model**

$$\begin{aligned}\frac{1}{4\rho^2}\Bigg[1 &- \frac{2}{\left[[\mu_b+\mu_c\gamma^{-1}\langle N\rangle]^2-4\sigma_c\sigma_g\rho\gamma^{-1}\langle v_N\rangle\mu_I\right]^{\frac{1}{2}}}\Bigg[\mu_b+\mu_c\gamma^{-1}\langle N\rangle \\ &- \frac{3\left[\mu_b+\mu_c\gamma^{-1}\langle N\rangle\right]\left[\sigma_c^2\gamma^{-1}\langle N^2\rangle+\sigma_b^2\right]}{2\left[[\mu_b+\mu_c\gamma^{-1}\langle N\rangle]^2-4\sigma_c\sigma_g\rho\gamma^{-1}\langle v_N\rangle\mu_I\right]}\Bigg] \\ &+ \frac{1}{[\mu_b+\mu_c\gamma^{-1}\langle N\rangle]^2-4\sigma_c\sigma_g\rho\gamma^{-1}\langle v_N\rangle\mu_I}\Bigg[\left[\mu_b+\mu_c\gamma^{-1}\langle N\rangle\right]^2+\sigma_c^2\gamma^{-1}\langle N^2\rangle+\sigma_b^2 \\ &- \frac{5\left[\mu_b+\mu_c\gamma^{-1}\langle N\rangle\right]^2\left[\sigma_c^2\gamma^{-1}\langle N^2\rangle+\sigma_b^2\right]}{[\mu_b+\mu_c\gamma^{-1}\langle N\rangle]^2-4\sigma_c\sigma_g\rho\gamma^{-1}\langle v_N\rangle\mu_I} \\ &- \frac{3\left[\sigma_c^2\gamma^{-1}\langle N^2\rangle+\sigma_b^2\right]^2}{[\mu_b+\mu_c\gamma^{-1}\langle N\rangle]^2-4\sigma_c\sigma_g\rho\gamma^{-1}\langle v_N\rangle\mu_I}\Bigg]\Bigg] \\ &< \phi_N\gamma^{-1}.\end{aligned} \tag{63}$$

#### A.6 Infeasibility condition

Next, we will investigate whether and when community dynamics become infeasible i.e., when a solution for the cavity self-consistency equations do not exist. From inspecting the self-consistency equations, we can see that the consumer and resource abundance distributions diverge to infinity when when $\langle \chi_R \rangle \to 0$ and $\langle v_N \rangle \to -\infty$. To determine when the system tends towards these limits, we will partially solve for $\langle v_N \rangle$ and $\langle \chi_R \rangle$. Plugging in eq. (34) to eq. (35) and rearranging yields the following expression:

$$\phi_N\gamma^{-1} = \frac{1}{2}\left[1-\frac{\mu_b+\mu_c\gamma^{-1}\langle N\rangle}{\sqrt{[\mu_b+\mu_c\gamma^{-1}\langle N\rangle]^2-4\sigma_c\sigma_g\rho\gamma^{-1}\langle v_N\rangle\mu_I}}\left[1-\frac{3\left[\sigma_c^2\gamma^{-1}\langle N^2\rangle+\sigma_b^2\right]}{2\left[[\mu_b+\mu_c\gamma^{-1}\langle N\rangle]^2-4\sigma_c\sigma_g\rho\gamma^{-1}\langle v_N\rangle\mu_I\right]}\right]\right]. \tag{64}$$

Since $\langle v_N \rangle \to -\infty$, $\langle N \rangle \to \infty$ and $\langle N^2 \rangle \to \infty$ in the infeasible limit, one can make a heuristic argument that $[\mu_b+\mu_c\gamma^{-1}\langle N\rangle] \div \sqrt{[\mu_b+\mu_c\gamma^{-1}\langle N\rangle]^2-4\sigma_c\sigma_g\rho\gamma^{-1}\langle v_N\rangle\mu_I} \to 0$ since the denominator contains two infinities whereas the numerator contains 1. Therefore, the r.h.s of eq. (64) tends to 1/2 as the system tends towards infeasibility. Therefore, $\phi_N\gamma^{-1}$ i.e., the consumer packing ratio can never exceed 1/2, consistent with [3].

### B Unified stability condition

We have thus far seen that the stability condition for the externally-supplied model, as well as the hybrid model, look rather different to the that of the self-renewing model [2, 4, 5]. To better understand the relationship between them, we now present an alternate derivation of the stability condition which will turn out to crucially rely on how resources respond to perturbations in their own depletion rates, i.e., on the statistics of the resource self-susceptibility $\chi_{00}$. We will also show that this alternate derivation leads to a unified stability condition that reduces to the known stability conditions for both models in some simple limits. Towards the end of this section, we also use the ideas developed here to discuss stability conditions for other classes of models with different resource dynamics, as well as reconcile apparent inconsistencies in the literature.

#### B.1 Sensitivity of resource abundances to perturbation

As above, we will perturb the surviving consumers as $N_i^+ \to N_i^+ + \varepsilon\eta_i$ and the resources as $R_\alpha \to R_\alpha + \varepsilon\eta_\alpha^{(R)}$. What distinguishes different models are their resource dynamics, and thus their resource sensitivities. We will first derive the stability condition for the externally-supplied resource model, paying close attention to the resource self-susceptibilities. As in the preceding calculation, for ease of notation, we will first collect the total resource depletion, both by dilution and by consumers, into the field $f$ as

$$\begin{aligned} f &= b_0 + \sum_i^S c_{i0} N_{i\backslash 0}^+ \\ &= \mu_b + \mu_c \gamma^{-1}\langle N\rangle + \frac{\sigma_c}{\sqrt{M}} \sum_{i=1}^{S^*} \left[\rho z_{g,i0} + \sqrt{1-\rho^2}\, z_{c,i0}\right] N_{i\backslash 0}^+ + \sigma_b z_{b,0}. \end{aligned} \tag{65}$$

Writing

$$F = \sigma_c \sigma_g \rho \gamma^{-1}\langle v_N\rangle \quad \text{and} \quad I_0 = \mu_I + \sigma_I z_{I,0}, \tag{66}$$

the cavity resource abundance may be written as the solution of a quadratic equation as

$$R_0 = \frac{f - \sqrt{f^2 - 4FI_0}}{2F}. \tag{67}$$

The dilution rate $b_0$ and the depletion generated by consumers enter $R_0$ through the same field $f$. By invoking the chain rule, we can characterize both responses using the diagonal resource susceptibility

$$\begin{aligned} \chi_{00} &= -\frac{\partial R_0}{\partial b_0} = -\frac{\partial R_0}{\partial f}\frac{\partial f}{\partial b_0} = -\frac{\partial R_0}{\partial f} \\ &= -\frac{1}{2F}\left[1 - \frac{f}{\sqrt{f^2 - 4FI_0}}\right] = \frac{R_0}{\sqrt{f^2 - 4FI_0}}. \end{aligned} \tag{68}$$

Note that this expression does not involve the binomial expansion and approximation used in the previous calculation. Averaging over the cavity ensemble results in the mean self-susceptibility we already used in the self-consistency equations previously, $\langle \chi_R\rangle = \langle \chi_{00}\rangle$.

We will now perturb the abundances of the surviving consumer species. This changes the local field $f$ and hence we can write

$$\frac{\mathrm{d}f}{\mathrm{d}\varepsilon} = \frac{\sigma_c}{\sqrt{M}} \sum_{i=1}^{S^*} \left[\rho z_{g,i0} + \sqrt{1-\rho^2}\, z_{c,i0}\right] \left[\frac{\mathrm{d}N_{i\backslash 0}^+}{\mathrm{d}\varepsilon} + \eta_i\right]. \tag{69}$$

Since the consumers affect $R_0$ only through $f$, we can use the chain rule to get an expression for how the resource abundance responds to perturbations in the consumer abundances

$$\frac{\mathrm{d}R_0}{\mathrm{d}\varepsilon} = \frac{\mathrm{d}R_0}{\mathrm{d}f}\frac{\mathrm{d}f}{\mathrm{d}\varepsilon} = -\chi_{00}\frac{\mathrm{d}f}{\mathrm{d}\varepsilon}. \tag{70}$$

Note again that Eqs. (68) and (70) do not involve the binomial expansion and approximation. To obtain the mean squared response, we now square Eq. (70) and average over cavity realizations. We will assume that in the large-$M$ limit, these two terms are weakly correlated, as in the previous calculation, leading to

$$\left\langle \left[\frac{\mathrm{d}R_0}{\mathrm{d}\varepsilon}\right]^2 \right\rangle = \left\langle \chi_{00}^2 \left[\frac{\mathrm{d}f}{\mathrm{d}\varepsilon}\right]^2 \right\rangle \approx \langle \chi_{00}^2 \rangle \left\langle \left[\frac{\mathrm{d}f}{\mathrm{d}\varepsilon}\right]^2 \right\rangle. \tag{71}$$

The first of these factors is the second moment of the resource self-susceptibility and will be a key quantity in the unified stability condition; we will come to it later. We next evaluate the second factor by squaring Eq. (69). This produces a double sum over pairs of surviving consumers:

$$\left\langle \left[\frac{\mathrm{d}f}{\mathrm{d}\varepsilon}\right]^2 \right\rangle = \frac{\sigma_c^2}{M}\sum_{i=1}^{S^*}\sum_{j=1}^{S^*}\left\langle \left[\rho z_{g,i0} + \sqrt{1-\rho^2}\, z_{c,i0}\right]\left[\rho z_{g,j0} + \sqrt{1-\rho^2}\, z_{c,j0}\right]\left[\frac{\mathrm{d}N^+_{i\backslash 0}}{\mathrm{d}\varepsilon} + \eta_i\right]\left[\frac{\mathrm{d}N^+_{j\backslash 0}}{\mathrm{d}\varepsilon} + \eta_j^{(N)}\right]\right\rangle, \tag{72}$$

which was calculated in eq. (48) to be approximately

$$\left\langle \left[\frac{\mathrm{d}f}{\mathrm{d}\varepsilon}\right]^2 \right\rangle \approx \sigma_c^2 \phi_N \gamma^{-1}\left[\left\langle \left[\frac{\mathrm{d}N_0^+}{\mathrm{d}\varepsilon}\right]^2 \right\rangle + 1\right]. \tag{73}$$

Substituting this result into eq. (71) gives

$$\left\langle \left[\frac{\mathrm{d}R_0}{\mathrm{d}\varepsilon}\right]^2 \right\rangle \approx \sigma_c^2 \phi_N \gamma^{-1}\langle \chi_{00}^2\rangle\left[\left\langle \left[\frac{\mathrm{d}N_0^+}{\mathrm{d}\varepsilon}\right]^2 \right\rangle + 1\right]. \tag{74}$$

Since all resources and consumers are statistically identical, $\langle \chi_{R,00}^2\rangle = \langle \chi_{R,\alpha\alpha}^2\rangle = \langle \chi_R^2\rangle$, meaning eq. (74) can be rewritten as

$$\left\langle \left[\frac{\mathrm{d}R_0}{\mathrm{d}\varepsilon}\right]^2 \right\rangle \approx \sigma_c^2 \phi_N \gamma^{-1}\langle \chi_R^2\rangle\left[\left\langle \left[\frac{\mathrm{d}N_0^+}{\mathrm{d}\varepsilon}\right]^2 \right\rangle + 1\right]. \tag{75}$$

This is an alternate compact form of the second moment of the resource response to perturbations in consumer abundances, analogous to Eq. (46). The factor $\phi_N\gamma^{-1} = S^*/M$ appears because only the $S^*$ surviving consumers contribute to the response of a resource. For another form of the resource supply dynamics, the expression for the resource self-susceptibility $\chi_{00}$ and its associated second moment $\langle \chi_R^2\rangle$ will change, but the form of Eq. (75) will typically remain the same whenever depletion by consumers enters through the same scalar field $f$ as resource dilution $b_0$. Note that for the self-renewing resource model, the depletion by consumers enters through changing the effective resource growth rate rather than the dilution rate, but mathematically, these conditions are equivalent since resource growth (or supply) rate in the self-renewing model enters as a linear term $\propto R_0$ in the resource supply dynamics [2].

#### B.2 Combined condition

We now combine the resource result with the consumer sensitivity derived in eq. (41). It tells us how a change in resource abundance changes a consumer. Equation (75) tells us how the resulting change in consumer abundance acts back on a resource. Stability is lost when these moments diverge and do not have a well-defined solution. Biologically, this corresponds to the case where perturbations in resources affect consumers, and those consumer changes feed back and generate a further change in resources. When this repeated response feedback becomes greatly amplified, the second moments diverge and the community becomes unstable.

To see this directly, we substitute eq. (75) into eq. (41):

$$\left\langle \left[\frac{\mathrm{d}N_0^+}{\mathrm{d}\varepsilon}\right]^2 \right\rangle \approx \frac{1}{\sigma_c^2 \rho^2 \langle \chi_R \rangle^2} \left[ \sigma_c^2 \phi_N \gamma^{-1} \langle \chi_R^2 \rangle \left[ \left\langle \left[\frac{\mathrm{d}N_0^+}{\mathrm{d}\varepsilon}\right]^2 \right\rangle + 1 \right] + 1 \right]. \tag{76}$$

Collecting the terms that contain the consumer sensitivity gives

$$\left[\rho^2 \langle \chi_R \rangle^2 - \phi_N \gamma^{-1} \langle \chi_R^2 \rangle\right] \left\langle \left[\frac{\mathrm{d}N_0^+}{\mathrm{d}\varepsilon}\right]^2 \right\rangle \approx \phi_N \gamma^{-1} \langle \chi_R^2 \rangle + \frac{1}{\sigma_c^2}. \tag{77}$$

Thus,

$$\left\langle \left[\frac{\mathrm{d}N_0^+}{\mathrm{d}\varepsilon}\right]^2 \right\rangle \approx \frac{\phi_N \gamma^{-1} \langle \chi_R^2 \rangle + 1/\sigma_c^2}{\rho^2 \langle \chi_R \rangle^2 - \phi_N \gamma^{-1} \langle \chi_R^2 \rangle}. \tag{78}$$

The resource sensitivity in eq. (75) diverges at the same point. Both sensitivities therefore remain finite and the community remains stable only when the denominator in eq. (78) is positive:

$$\rho^2 \langle \chi_R \rangle^2 > \phi_N \gamma^{-1} \langle \chi_R^2 \rangle. \tag{79}$$

Since $\phi_N \gamma^{-1} = S^*/M$, the stability condition becomes

**Unified stability condition**

$$\rho^2 > \frac{S^*}{M} \frac{\langle \chi_R^2 \rangle}{\langle \chi_R \rangle^2}. \tag{80}$$

Note that this boxed stability condition applies to both the externally-supplied and self-renewing resource models, due to arguments we have made previously. This stability condition has a rather simple interpretation. The packing fraction $S^*/M$ counts how many surviving consumers contribute to the response of each resource, and is a standard and known determinant of ecological stability. The mean susceptibility $\langle \chi_R \rangle = \langle \chi_{00} \rangle$ enters when a resource perturbation changes the consumers, whereas the second moment $\langle \chi_R^2 \rangle$ enters when the consumer response acts back on the resources. Their normalized ratio therefore analytically describes how including explicit resource dynamics changes the stability condition for complex ecological communities.

#### B.3 Relation to stability condition for self-renewing resources

We can also show that the same unified stability condition applies to a resource model with self-renewing resources. Without external resource influx, resources can become extinct. We perturb only the $M^*$ surviving resources,

$R^+_\alpha \to R^+_\alpha + \varepsilon\eta^{(R)}_\alpha$, and write their survival fraction as $\phi_R = M^*/M$. As in the rest of the linear-response calculation, we assume that the perturbation is small and does not change the set of surviving resources. Because the sum in the consumer response now contains $M^*$ rather than $M$ terms, the factor $\phi_R$ multiplies both the induced response of a surviving resource and its direct perturbation. The consumer sensitivity is therefore

$$\left\langle \left[\frac{\mathrm{d}N^+_0}{\mathrm{d}\varepsilon}\right]^2 \right\rangle = \frac{\phi_R}{\sigma_c^2\rho^2\langle\chi_R\rangle^2}\left[\left\langle \left[\frac{\mathrm{d}R^+_0}{\mathrm{d}\varepsilon}\right]^2 \right\rangle + 1\right]. \tag{81}$$

The response of resources to perturbations in the abundances of surviving consumers remains similar to Eq. (75), albeit we again restrict it to the set of surviving resources, which gives us

$$\left\langle \left[\frac{\mathrm{d}R^+_0}{\mathrm{d}\varepsilon}\right]^2 \right\rangle = \sigma_c^2\phi_N\gamma^{-1}\langle\chi_R^2\rangle^+ \left[\left\langle \left[\frac{\mathrm{d}N^+_0}{\mathrm{d}\varepsilon}\right]^2 \right\rangle + 1\right], \tag{82}$$

where $\langle\chi_R^2\rangle^+$ is the analog of the second moment of the resource self-susceptibility in Eq. (80), except again restricted to the surviving resources. Note that extinct resources have zero susceptibility. Using this, we obtain the following relation between the first and second moments of the resource self-susceptibility, which we will use later:

$$\langle\chi_R\rangle = \phi_R\langle\chi_R\rangle^+. \tag{83}$$

$$\langle\chi_R^2\rangle = \phi_R\langle\chi_R^2\rangle^+. \tag{84}$$

Substituting Eq. (82) into Eq. (81), we get

$$\left\langle \left[\frac{\mathrm{d}N^+_0}{\mathrm{d}\varepsilon}\right]^2 \right\rangle = \frac{\phi_R}{\sigma_c^2\rho^2\langle\chi_R\rangle^2}\left[\sigma_c^2\phi_N\gamma^{-1}\langle\chi_R^2\rangle^+\left[\left\langle \left[\frac{\mathrm{d}N^+_0}{\mathrm{d}\varepsilon}\right]^2 \right\rangle + 1\right] + 1\right]. \tag{85}$$

Solving for the consumer sensitivity, we get

$$\left\langle \left[\frac{\mathrm{d}N^+_0}{\mathrm{d}\varepsilon}\right]^2 \right\rangle = \frac{\phi_R\phi_N\gamma^{-1}\langle\chi_R^2\rangle^+ + \phi_R/\sigma_c^2}{\rho^2\langle\chi_R\rangle^2 - \phi_R\phi_N\gamma^{-1}\langle\chi_R^2\rangle^+}. \tag{86}$$

Substituting our expression for $\langle\chi_R^2\rangle^+$ in Eq. (84) yields

$$\left\langle \left[\frac{\mathrm{d}N^+_0}{\mathrm{d}\varepsilon}\right]^2 \right\rangle = \frac{\phi_N\gamma^{-1}\langle\chi_R^2\rangle + \phi_R/\sigma_c^2}{\rho^2\langle\chi_R\rangle^2 - \phi_N\gamma^{-1}\langle\chi_R^2\rangle}. \tag{87}$$

the denominator in Eq. (87) is the same as the denominator in Eq. (78). This shows that point at which the response response moments diverge and the community becomes unstable is therefore still determined by the boxed unified condition (80).

To simplify this further, we take the simple and typical case where all resources have the same intrinsic growth rate $b_\alpha$ and carrying capacity $K_\alpha$. In this limit, we may assume that all surviving resources have a common self-susceptibility $\chi_{\text{common}}$, while extinct resources have zero self-susceptibility. Note that in this model, the self-susceptibilities are usually computed due to changes in the resource growth rate rather than their outflux rate. However, this will only change the overall sign of the self-susceptibility $\chi_{\text{common}}$, which will not affect the stability condition since it only involves the square of the self-susceptibility and its second moment. Using this simplification, we can see that the $\langle \chi_R \rangle$ in eq. (83) can be written as

$$\langle \chi_R \rangle = \phi_R \chi_{\text{common}}, \tag{88}$$

while $\langle \chi_R^2 \rangle$ from eq. (84) can be written as

$$\langle \chi_R^2 \rangle = \phi_R \chi_{\text{common}}^2. \tag{89}$$

Thus, this simplified limit, we see that the unified stability condition reduces to

$$\rho^2 > \frac{S^*}{M} \frac{\phi_R \chi_{\text{common}}^2}{[\phi_R \chi_{\text{common}}]^2} = \frac{S^*}{M} \frac{1}{\phi_R} \tag{90}$$

Substituting $\phi_R = M^*/M$ yields the known stability condition for the self-renewing model [2]:

$$\rho^2 > \frac{S^*}{M*} \tag{91}$$

Our unified stability condition further emphasizes the susceptibility of resources at or near extinction as a primary driver of stability or instability. In the self-renewing model, the absence of external resource influx permits resource extinctions. For extinct resources, the self-susceptibility is zero, while the surviving resources respond with susceptibility $\chi_{\text{common}}$ in the limit considered above. The first and second moments then contain different powers of the resource survival fraction, so their ratio simplifies to $M/M^*$ and gives $\rho^2 > S^*/M^*$. In the externally-supplied model, positive resource influx prevents resources from going extinct. The resource susceptibilities therefore cannot be simply separated into surviving and extinct parts, and the second moment of the self-susceptibility must instead be evaluated over the full distribution of resources. This is why resources with large depletion rate susceptibilities can have a disproportionate effect on the stability boundary. The presence or absence of resource influx changes this susceptibility distribution, and hence distinguishes the stability conditions of the two resource models.

#### B.4 Relation to stability condition for externally supplied resources

To clarify the connection between the two stability conditions derived here and in the previous section, consider the expression in Eq. (68). Squaring and taking its expectation gives us

$$\begin{aligned} \langle \chi_{00}^2 \rangle &= \left\langle \left[ -\frac{1}{2F} \left[ 1 - \frac{f}{\sqrt{f^2 - 4FI_0}} \right] \right]^2 \right\rangle \\ \Rightarrow 4F^2 \langle \chi_R^2 \rangle &= \left\langle \left[ 1 - \frac{f}{\sqrt{f^2 - 4FI_0}} \right]^2 \right\rangle . \end{aligned} \tag{92}$$

This equation is directly related to the long bracketed term in Eq. (63). To see this, note that before evaluating the long bracketed term in Eq. (63) using binomial expansion, the stability condition is

$$\frac{1}{4\rho^2} \left\langle \left[ 1 - \frac{f}{\sqrt{f^2 - 4FI_0}} \right]^2 \right\rangle < \phi_N \gamma^{-1}. \tag{93}$$

Substituting eq. (92) into this expression gives

$$\frac{F^2 \langle \chi_R^2 \rangle}{\rho^2} < \phi_N \gamma^{-1}. \tag{94}$$

Substituting in $v_N$ from (34) to $F$ in (66) yields $F^2 = [\phi_N \gamma^{-1}]^2 / \langle \chi_R^2 \rangle$. Finally, we can substitute this expression for $F$ into eq. (94), which gives

$$\frac{[\phi_N \gamma^{-1}]^2 \langle \chi_R^2 \rangle}{\rho^2 \langle \chi_R \rangle^2} < \phi_N \gamma^{-1} \tag{95}$$

$$\Rightarrow \rho^2 > \phi_N \gamma^{-1} \frac{\langle \chi_R^2 \rangle}{\langle \chi_R \rangle^2} = \frac{S^*}{M} \frac{\langle \chi_R^2 \rangle}{\langle \chi_R \rangle^2}, \tag{96}$$

which is the unified stability condition in the box Eq. (80). The unified condition is useful when the susceptibility moment can be evaluated directly, or when we wish to compare models with different local resource dynamics. For example, if we take the simple limit where all resources have the same susceptibility $\chi_{\text{common}}$, then the condition reduces to

$$\rho^2 > \frac{S^*}{M}, \tag{97}$$

which is comparable to the condition for self-renewing resources.

However, if we want to evaluate the susceptibilities and obtain a general stability condition for externally-supplied resources, we can plug the expression for $F$ into eq. (92), rearrange the expression to make $\langle \chi_R^2 \rangle$ the subject, then plug this expression into eq. (96) to yield

$$\rho^2 > \frac{M}{4S^*} \left\langle \left[ 1 - \frac{b_\alpha^{\text{eff}}}{\sqrt{(b_\alpha^{\text{eff}})^2 + \frac{4S^* I_\alpha}{M \langle \chi_R \rangle}}} \right]^2 \right\rangle, \tag{98}$$

.

This alternative derivation leaves the same average in terms of $\langle \chi_R^2 \rangle$ rather than fully evaluating the bracket. If we want an explicit approximation in terms of the order parameters of the externally-supplied model, we can perform a first order binomial expansion on the bracket. This yields the original stability condition.

#### B.5 Generalisation to other forms of resource dynamics

This alternate calculation also suggests how to treat other local resource dynamics. For each model, one first determines the response of a single resource and then computes the first and second moments of that response over the self-consistent resource distribution. We hypothesize that models where resources can go extinct will have a stability condition similar to the self-renewing limit, especially when resources are supplied at the same rate. In contrast, models where resources are externally supplied and cannot go extinct will have a stability condition similar to the general unified stability condition, albeit with a different form of $\langle \chi_R^2 \rangle$. The hybrid model considered above provides one example that interpolates between the two limits.

More complex resource dynamics, where the influx rates depend on the current state of the community might lead to a different resource self-susceptibility distribution $\chi_{00}$ and therefore have a different stability boundary. Cross-feeding of metabolic byproducts could be an example of such complex resource dynamics. Further, if there were direct resource-resource interactions, we might need a slightly more general distribution over the full susceptibility matrix, rather than just the diagonal elements.

This perspective also explains what is lost when models neglect resource dynamics and replace them with effective consumer–consumer interactions. In such descriptions, the resource susceptibility factor is usually fixed by hand, absorbed into the effective interactions, or approximated by a single value. Thus, it is no longer amenable to feedback from consumer dynamics, and thus might lead to incorrect estimates of the stability boundary. Keeping the resource dynamics explicit allows this factor to emerge naturally from community feedbacks, resulting in a more accurate description of the stability boundary. It also cleanly decomposes how different factors such as consumer diversity, resource diversity, and resource susceptibilities impact community stability.

### C Hybrid model with both externally-supplied and self-renewing resource supply

In this section, we apply the unified calculation above to a model with both external resource influx and resource self-renewal. We first derive its stability condition when the influx is positive, and then show how the condition reduces to the self-renewing result when the influx is zero.

#### C.1 Model set up

$$\begin{aligned}\frac{dR_\alpha}{dt} &= \underbrace{I_\alpha + R_\alpha\left[b_\alpha - \frac{R_\alpha}{K_\alpha}\right]}_{h_\alpha(R_\alpha)} - R_\alpha \sum_{i=1}^{S} c_{i\alpha} N_i. \\ \frac{dN_i}{dt} &= N_i\left[\sum_{\alpha=1}^{M} g_{i\alpha} R_\alpha - d_i\right].\end{aligned} \tag{99}$$

$I_\alpha$ is the influx rate and $b_\alpha$ is the intrinsic resource growth rate. The latter can be interpreted as the net biological growth rate minus the outflux rate, and can therefore be positive or negative. $K_\alpha$ is the carrying capacity of resource $\alpha$ and controls the strength of resource self-inhibition. In the calculation below, we set $K_\alpha = 1$.

#### C.2 Cavity consumer and resource abundance distributions

The equation for the cavity consumer remains unchanged from the externally-supplied model. The cavity resource $R_0$, however, satisfies

$$0 = \mu_I + \sigma_I z_{I,0} + R_0\left[\mu_b - \mu_c\gamma^{-1}\langle N\rangle + \sqrt{\sigma_c^2\gamma^{-1}\langle N^2\rangle + \sigma_b^2} Z_R\right] + R_0^2[\sigma_g\sigma_c\rho\gamma^{-1}\langle v_N\rangle - 1]. \tag{100}$$

Solving this equation gives the following expression for the cavity resource abundance:

$$\begin{aligned}R_0 = &-\frac{\mu_b - \mu_c\gamma^{-1}\langle N\rangle + \sqrt{\sigma_c^2\gamma^{-1}\langle N^2\rangle + \sigma_b^2} Z_R}{2\left[\sigma_c\sigma_g\rho\gamma^{-1}\langle v_N\rangle - 1\right]} \\ &- \frac{\sqrt{[\mu_b - \mu_c\gamma^{-1}\langle N\rangle + \sqrt{\sigma_c^2\gamma^{-1}\langle N^2\rangle + \sigma_b^2} Z_R]^2 - 4[\sigma_c\sigma_g\rho\gamma^{-1}\langle v_N\rangle - 1][\mu_I + \sigma_I z_{I,0}]}}{2\left[\sigma_c\sigma_g\rho\gamma^{-1}\langle v_N\rangle - 1\right]}.\end{aligned} \tag{101}$$

#### C.3 Stability condition

As above, we perturb the surviving consumers and the resources and calculate the second moments of their responses. When $I_\alpha > 0$ for every resource, resources cannot go extinct. Thus $M^* = M$ and $\phi_R = 1$, and the consumer sensitivity is given by Eq. (41). When the influx is zero, resources can go extinct. We then perturb only the $M^*$ surviving resources and use the consumer sensitivity in Eq. (81).

We first consider $I_\alpha > 0$. For ease of notation, we write $a_0$ as

$$a_0 = \mu_b - \mu_c\gamma^{-1}\langle N\rangle + \sqrt{\sigma_c^2\gamma^{-1}\langle N^2\rangle + \sigma_b^2}\, Z_R. \tag{102}$$

We will also use

$$H = \sigma_g\sigma_c\rho\gamma^{-1}\langle v_N\rangle - 1, \tag{103}$$

and

$$I_0 = \mu_I + \sigma_I z_{I,0}. \tag{104}$$

Using this notation, Eq. (101) becomes

$$R_0 = \frac{-a_0 - \sqrt{a_0^2 - 4HI_0}}{2H}. \tag{105}$$

The intrinsic growth rate and the depletion generated by consumers enter $R_0$ through $a_0$ with opposite signs. An increase in $a_0$ is therefore equivalent to a decrease in the resource depletion rate. We can characterize this response using the resource self-susceptibility

$$\chi_{R,00} = \frac{\partial R_0}{\partial a_0} = -\frac{1}{2H}\left[1 + \frac{a_0}{\sqrt{a_0^2 - 4HI_0}}\right]. \tag{106}$$

Averaging over resources gives the mean susceptibility

$$\langle \chi_R\rangle = \langle \chi_{R,00}\rangle\,. \tag{107}$$

Perturbing consumer abundances changes $a_0$ through the consumer depletion term. Multiplying this change by $\partial R_0/\partial a_0$, squaring the result, and averaging over resources, we get

$$\left\langle\left[\frac{\mathrm{d}R_0}{\mathrm{d}\varepsilon}\right]^2\right\rangle \approx \sigma_c^2\phi_N\gamma^{-1}\left\langle\chi_{R,00}^2\right\rangle\left[\left\langle\left[\frac{\mathrm{d}N_0^+}{\mathrm{d}\varepsilon}\right]^2\right\rangle + 1\right] = \sigma_c^2\phi_N\gamma^{-1}\left\langle\chi_R^2\right\rangle\left[\left\langle\left[\frac{\mathrm{d}N_0^+}{\mathrm{d}\varepsilon}\right]^2\right\rangle + 1\right]. \tag{108}$$

Substituting Eq. (108) into the consumer sensitivity in Eq. (41), and requiring the resulting second moments to remain finite, gives us the unified stability condition:

#### Hybrid model stability condition

$$\rho^2 > \frac{S^*}{M}\frac{\langle \chi_R^2 \rangle}{\langle \chi_R \rangle^2}. \tag{109}$$

To connect this condition to one that contains a more explicit expression for $\langle \chi_R^2 \rangle$, we can instead square Eq. (106) and average over resources, which gives

$$\left\langle \left[ 1 + \frac{a_0}{\sqrt{a_0^2 - 4HI_0}} \right]^2 \right\rangle = 4H^2 \left\langle \chi_R^2 \right\rangle. \tag{110}$$

Using the same binomial expansion as in the externally-supplied calculation to evaluate these averages, Eq. (108) becomes

$$\begin{aligned}
\left\langle \left[ \frac{\mathrm{d}R_0}{\mathrm{d}\varepsilon} \right]^2 \right\rangle \approx{}& \frac{\sigma_c^2 \phi_N \gamma^{-1}}{4\left[\sigma_g \sigma_c \rho \gamma^{-1} \langle v_N \rangle - 1\right]^2} \left[ \left\langle \left[ \frac{\mathrm{d}N_0^+}{\mathrm{d}\varepsilon} \right]^2 \right\rangle + 1 \right] \\
&\times \left[ 1 + \frac{2}{\left[ \left[\mu_b - \mu_c \gamma^{-1} \langle N \rangle\right]^2 - 4\left[\sigma_g \sigma_c \rho \gamma^{-1} \langle v_N \rangle - 1\right] \mu_I \right]^{\frac{1}{2}}} \left[ \mu_b - \mu_c \gamma^{-1} \langle N \rangle \right. \right. \\
&\qquad \left. - \frac{3\left[\mu_b - \mu_c \gamma^{-1} \langle N \rangle\right] \left[\sigma_c^2 \gamma^{-1} \langle N^2 \rangle + \sigma_b^2\right]}{2\left[ \left[\mu_b - \mu_c \gamma^{-1} \langle N \rangle\right]^2 - 4\left[\sigma_g \sigma_c \rho \gamma^{-1} \langle v_N \rangle - 1\right] \mu_I \right]} \right] \\
&+ \frac{1}{\left[\mu_b - \mu_c \gamma^{-1} \langle N \rangle\right]^2 - 4\left[\sigma_g \sigma_c \rho \gamma^{-1} \langle v_N \rangle - 1\right] \mu_I} \left[ \left[\mu_b - \mu_c \gamma^{-1} \langle N \rangle\right]^2 + \sigma_c^2 \gamma^{-1} \langle N^2 \rangle + \sigma_b^2 \right. \\
&\qquad - \frac{5\left[\mu_b - \mu_c \gamma^{-1} \langle N \rangle\right]^2 \left[\sigma_c^2 \gamma^{-1} \langle N^2 \rangle + \sigma_b^2\right]}{\left[\mu_b - \mu_c \gamma^{-1} \langle N \rangle\right]^2 - 4\left[\sigma_g \sigma_c \rho \gamma^{-1} \langle v_N \rangle - 1\right] \mu_I} \\
&\qquad \left. \left. - \frac{3\left[\sigma_c^2 \gamma^{-1} \langle N^2 \rangle\right]^2 + 6\sigma_b^2 \left[\sigma_c^2 \gamma^{-1} \langle N^2 \rangle\right] + 3\sigma_b^4}{\left[\mu_b - \mu_c \gamma^{-1} \langle N \rangle\right]^2 - 4\left[\sigma_g \sigma_c \rho \gamma^{-1} \langle v_N \rangle - 1\right] \mu_I} \right] \right].
\end{aligned} \tag{111}$$

Thus, Eq. (111) is an approximate evaluation of the same second moment $\langle \chi_R^2 \rangle$ that appears in the boxed condition. This condition can be evaluated using quantities that we have already determined through the self-consistency equations, whereas the boxed condition is more compact, but does not contain an explicit method to evaluate $\langle \chi_R^2 \rangle$.

To look at this expression in various limits, we may set the resource influx to zero. Resources may then either go extinct or survive. Setting $\mu_I = \sigma_I = 0$ and following similar steps as in the previous sections, we can show that

$$\frac{\langle \chi_R^2 \rangle}{\langle \chi_R \rangle^2} = \frac{1}{\phi_R} = \frac{M}{M^*}. \tag{112}$$

Substituting this result into Eq. (109) gives $\rho^2 > S^*/M^*$, which is the self-renewing stability condition in Eq. (91). Thus, in the limit of no resource influx where resources can go extinct, we obtain the same stability condition in the hybrid model as in the self-renewing model.

We may also consider two other limits in the presence of resource influx, i.e., $I_\alpha > 0$, so that all resources survive and $M^* = M$. First, disallowing resources to get diluted by setting $b_\alpha = 0$ reduces Eq. (111) to

$$\left\langle \left[\frac{\mathrm{d}R_0}{\mathrm{d}\varepsilon}\right]^2 \right\rangle \approx \frac{\sigma_c^2 \phi_N \gamma^{-1}}{4\left[\sigma_g\sigma_c\rho\gamma^{-1}\langle v_N\rangle - 1\right]^2} \left[\left\langle \left[\frac{\mathrm{d}N_0^+}{\mathrm{d}\varepsilon}\right]^2 \right\rangle + 1\right]$$
$$\times \Bigg[1 - \frac{2}{\left[\left[\mu_c\gamma^{-1}\langle N\rangle\right]^2 - 4\left[\sigma_g\sigma_c\rho\gamma^{-1}\langle v_N\rangle - 1\right]\mu_I\right]^{\frac{1}{2}}} \Bigg[\mu_c\gamma^{-1}\langle N\rangle - \frac{3\left[\mu_c\gamma^{-1}\langle N\rangle\right]\left[\sigma_c^2\gamma^{-1}\langle N^2\rangle\right]}{2\left[\left[\mu_c\gamma^{-1}\langle N\rangle\right]^2 - 4\left[\sigma_g\sigma_c\rho\gamma^{-1}\langle v_N\rangle - 1\right]\mu_I\right]}\Bigg]$$
$$+ \frac{1}{\left[\mu_c\gamma^{-1}\langle N\rangle\right]^2 - 4\left[\sigma_g\sigma_c\rho\gamma^{-1}\langle v_N\rangle - 1\right]\mu_I} \Bigg[\left[\mu_c\gamma^{-1}\langle N\rangle\right]^2 + \sigma_c^2\gamma^{-1}\langle N^2\rangle - \frac{5\left[\mu_c\gamma^{-1}\langle N\rangle\right]^2\left[\sigma_c^2\gamma^{-1}\langle N^2\rangle\right]}{\left[\mu_c\gamma^{-1}\langle N\rangle\right]^2 - 4\left[\sigma_g\sigma_c\rho\gamma^{-1}\langle v_N\rangle - 1\right]\mu_I} - \frac{3\left[\sigma_c^2\gamma^{-1}\langle N^2\rangle\right]^2}{\left[\mu_c\gamma^{-1}\langle N\rangle\right]^2 - 4\left[\sigma_g\sigma_c\rho\gamma^{-1}\langle v_N\rangle - 1\right]\mu_I}\Bigg]\Bigg]. \tag{113}$$

This has a similar structure to the stability condition for the externally supplied model in Eq. (63), and shows that the presence of resource influx alone is sufficient to result in a stability condition that is rather different from the self-renewing resource model.

Second, we can also remove resource self-inhibition by setting $K_\alpha \to \infty$. A finite steady-state resource abundance then requires $b_\alpha < 0$. We can thus write the corresponding resource growth rate as instead a dilution rate by setting $\tilde{b}_\alpha = -b_\alpha$ and denote its mean by $\mu_b$. With these substitutions, the resource response reduces to that of the externally-supplied model in Eq. (77). Thus, removing self-inhibition while retaining positive resource influx recovers the externally-supplied model, whereas removing influx recovers the self-renewing model. In both cases, the resource dynamics are an integral part of the stability condition, which depends on the first and second moments of the resource self-susceptibility $\chi_{00}$.

### D Numerical details

All codes for running simulations and solving self-consistency equations can be found in this repository: `https://github.com/JamilaRowlandChandler/CRM-Resource-supply-vs-Stability`. This repository also contains notebooks with examples for running simulations and solving self-consistency equations.

#### D.1 Simulations

The models used in simulations are of the form

$$\frac{\mathrm{d}R_\alpha}{\mathrm{d}t} = h_\alpha(R_\alpha) - R_\alpha \sum_{i=1}^{S} c_{i\alpha} N_i + \zeta, \quad \frac{\mathrm{d}N_i}{\mathrm{d}t} = N_i \left[\sum_{\alpha=1}^{M} g_{i\alpha} R_\alpha - d_i\right] + \zeta, \tag{114}$$

where the resource supply function $h_\alpha(R_\alpha)$ took the form

$$\textbf{Self-renewing}: R_\alpha \left[b_\alpha - \frac{R_\alpha}{K_\alpha}\right]$$
$$\textbf{Externally-supplied}: I_\alpha - b_\alpha R_\alpha \tag{115}$$
$$\textbf{Hybrid supply}: I_\alpha + R_\alpha \left[b_\alpha - \frac{R_\alpha}{K_\alpha}\right]$$

Parameters were sampled from the following distributions:

$$c_{i\alpha} = \frac{\mu}{M} + \frac{\sigma}{\sqrt{M}}\left[\rho z_{g,i\alpha} + \sqrt{1-\rho^2} z_{c,i\alpha}\right], \quad g_{i\alpha} = \frac{\mu}{M} + \frac{\sigma}{\sqrt{M}} z_{g,i\alpha}, \quad I_\alpha = I, \quad b_\alpha = 1, \quad K_\alpha = K, \quad d_i = 1, \quad \zeta = 10^{-8}, \tag{116}$$

where $\zeta$ is the "migration rate". This term is added to reduce numerical instability in our simulations, which occurs when some species and resource abundances are very small but not extinct. We neglect it in our analytical calculations because it is very small.

We generated communities and simulated their dynamics in Python using *numpy* and scipy's ODE solver solve_ivp, and in Mathematica using *NDSolve*. For each set of parameter distributions, 20 communities were sampled.

Simulation data in fig. 2 and 3 in the main text was generated using Python. The parameters of each community were sampled from their respective normal distributions using *numpy.random.randn*. Community dynamics were then simulated from 1 set of initial abundances sampled from Uniform(min. $= \zeta$, max. $= 2/M$). Dynamics were simulated for $t = 1000$ using the *LSODA* routine from scipy.integrate.solve_ivp, which is built for efficiently dealing with stiff (and non-stiff) ODE problems. An *unbounded growth* condition was also included to terminate the simulation when abundances grew unbounded. Hyper-parameters: the solver's relative tolerance was set to $10^{-7}$, absolute tolerance to $10^{-9}$. Dynamics were saved at d$t = 5$. Simulation data in fig. S2 was simulated in much the same way, except $t = 7000$.

Simulation data in fig. S1 was generated using Mathematica. The parameters of each community were sampled from their respective normal distributions using NormalDistribution. Community dynamics were then simulated from 1 set of initial abundances sampled from Uniform(min. $= \zeta$, max. $= 2/M$). Dynamics were estimated using the *StiffnessSwitching* routine with the *ExplicitRungeKutta* method from NDSolve. An *unbounded growth* condition was also included to terminate the simulation when abundances grew unbounded. Hyper-parameters: the solver's relative tolerance was set to $10^{-7}$, absolute tolerance to $10^{-9}$. Dynamics were evaluated until $t = 400$ at every d$t = 2$ timestep.

**Figure 2** – For both models, $\rho$ varied between 0.1 and 1.0 in increments of 0.1, $\sigma$ varied between 2.0 and 12.0 in increments of 0.5. Other shared parameters were fixed to $M = S = 150$, $\mu = 50.0$, $\mu_d = 1.0$, $\sigma_d = 0.0$, $\mu_b = 1.0$, $\sigma_b = 0,0$. In the self-renewing model (Fig. 2A), $K = 1.0$. In the externally-supplied model (Fig. 2B), $\mu_I = 1.0$, $\sigma_I = 0.0$.

**Figure 3A** – From left to right, external resource influx and self-inhibition varied as follows: $\mu_I = 10^{-5}$, $\sigma_I = 0.0$, $K = 1/10^{-5}$; $\mu_I = 10^{-5}$, $\sigma_I = 0.0$, $K = 1$; $\mu_I = 1$, $\sigma_I = 0.0$, $K = 1/10^{-5}$; $\mu_I = 1$, $\sigma_I = 0.0$, $K = 1$. Within each plot, $\rho$ varied between 0.1 and 1.0 in increments of 0.1, $\sigma$ varied between 2.0 and 12.0 in increments of 0.5. All other parameters were fixed to $M = S = 150$, $\mu = 50.0$, $\mu_d = 1.0$, $\sigma_d = 0.0$, $\mu_b = 1.0$, $\sigma_b = 0.0$.

**Figure 3B** – $\rho$ varied between 0.1 and 1.0 in increments of 0.1, $I_\alpha$ varied between $10^{-5}$ and $10^0$, where the exponent varied in increments of 0.5. All other parameters were fixed to $M = S = 150$, $\mu = 50.0$, $\sigma = 4.0$, $\mu_d = 1.0$, $\sigma_d = 0.0$, $\mu_b = 1.0$, $\sigma_b = 0.0$, $K = 1$.

**Figure 3C** – Simulations were run across the boundary between stability and chaos for $\rho$ and $I_\alpha$. Values of $I_\alpha$ were the same as Fig. 3B. For each value of $I_\alpha$, a corresponding value of $\rho$ was selected along the phase boundary. The values of $\rho$ selected were $[0.85, 0.85, 0.8, 0.75, 0.7, 0.55, 0.4]$. All other parameters were the same as Fig. 3B. For each set of parameter distributions, 80 communities were sampled.

**Figure S1A** $\rho$ varied between 0.1 and 1.0 in increments of 0.1, $\sigma$ varied between 1.0 and 2.0 in increments of 0.1. All other parameters were fixed to $M = S = 150$, $\mu = 1.0$, $\mu_d = 1.0$, $\sigma_d = 0.1$, $\mu_I = 1.0$, $\sigma_I = 0.1$, $\mu_b = 1.0$, $\sigma_b = 0.0$.

**Figure S2B** $I_\alpha$ was varied between $10^{-8}$ and $10^{-0.5}$. All other parameters were fixed to $M = S = 150$, $\rho = 0.4$, $\mu = 50.0$, $\sigma = 10.5$, $\mu_d = 1.0$, $\sigma_d = 0.0$, $\mu_b = 1.0$, $\sigma_b = 0.0$.

#### D.2 Estimating community stability

We determined the stability of each community by numerically estimating its maximum Lyapunov exponent. In a continuous-time system, we can define the maximum Lyapunov exponent as follows [6]. Let's say we have two nearby trajectories of species and resource abundances $\mathbf{x}(t)$ in phase space. The first trajectory is denoted as $\mathbf{x}(t)$ and the nearby trajectory is denoted as $\mathbf{x}(t) + \boldsymbol{\delta}(t)$, where $\boldsymbol{\delta}(0)$ is the initial distance between the trajectories. In the limit of the initial distance $\boldsymbol{\delta}(0)$ going to zero and the time for which the two trajectories evolve going to $\infty$, we can define the maximum Lyapunov exponent $\lambda_{\max}$ in terms of the long-term distance $\boldsymbol{\delta}(t)$ between the trajectories as:

$$\lambda_{\max} = \lim_{\substack{\delta(0)\to 0 \\ t\to\infty}} \frac{1}{t} \log\left[\frac{|\boldsymbol{\delta}(t)|}{|\boldsymbol{\delta}(0)|}\right] \tag{117}$$

We can see that if $\lambda_{\max} < 0$, the absolute distance between trajectories $|\delta(t)|$ decays over time, indicating the system converges to the same steady state. If $\lambda_{\max} > 0$, the trajectories diverge over time, indicating the system is unstable. We use this definition to calculate each community's maximum Lyapunov exponent using the algorithm from [7], summarised by Hrothgar (2015) and Koehler (2024).

##### Algorithm for estimating $\lambda_{\max}$

1. For the CRM, extract the final species and resource abundances from the end of simulations (detailed in Appendix D.1). This is the "original trajectory" at $t = 0$.
2. Initialise a "perturbed trajectory". Perturb the original abundances by some small amount $\boldsymbol{\delta}(0)$ i.e., the Euclidean distance between the original and perturbed trajectory is $\boldsymbol{\delta}(0)$.
   - We set $\boldsymbol{\delta}(0)$ to $10^{-6}$.
3. Simulate the dynamics of the original and perturbed trajectory, computing the normalised (log) Euclidean distance between them $\log(\boldsymbol{\delta}(t))$ at each time step. Continue to simulate dynamics until $\log(\boldsymbol{\delta}(t))$ becomes approximately constant within some tolerance.
   - We found simulating dynamics for $t = 1000$ was sufficient to achieve this.
4. To estimate the maximum Lyapunov exponent $\lambda_{\max}$, fit a line to the log distance between trajectories $\log(\boldsymbol{\delta}(t))$ in the region where it varies with time. The best-fit slope is the estimated maximum Lyapunov exponent $\lambda_{\max}$.

#### D.3 Estimating community feasibility

We observed that communities with infeasible dynamics often exhibited unbounded-like growth or were too numerically unstable for the ODE solver to handle. These dynamics stopped the ODE solver before its set end time $t$ because it either activated the unbounded_growth event function, which stops simulations when any consumer or resource grows above $10^6$, or terminated of its own accord due to numerical instability. Therefore, we determined a community was feasible if it ran until the end time, and infeasible if it terminated at an earlier time step.

#### D.4 Estimating the packing ratio

We defined the number of surviving consumers (resources) as the number of consumers (resources) with abundances greater than extinction threshold $e$ at the end of simulations. We arbitrarily set $e$ to $10^{-4}$. These quantities were then used to estimate the packing ratio.

#### D.5 Estimating the sensitivity of resources to changes in depletion

To estimate the sensitivity of resources to changes in their total depletion rate $(\mathrm{d}R_\alpha/\mathrm{d}C_\alpha)^2$, as plotted in main text Fig. 3C i, we first sorted resource abundances $R_\alpha$ in order of their total depletion rates $C_\alpha = \sum_i^S c_{i,\alpha} N_i$. We then calculated the gradient of $R_\alpha$ in terms of $C_\alpha$ using *numpy.gradient* to estimate $\mathrm{d}R_\alpha/\mathrm{d}C_\alpha$. This was then squared to obtain the $(\mathrm{d}R_\alpha/\mathrm{d}C_\alpha)^2$, then binned by their average using *scipy.stats.binned.statistic*. Hyper-parameters: the number of bins *bins* was ad-hoc set to 15.

To determine whether sensitivity of near-extinction resources to changes in their depletion affected boom-bust dynamics, we needed to estimate their maximum sensitivity. These are plotted in Fig. 3C ii in the main text. As shown in Fig. 3C i in the main text, resources with a total depletion rate $C_\alpha$ that is approximately equal to the intrinsic resource growth rate $b_\alpha = 1$ are near extinction. In the chaotic phase, these resources have a spike in sensitivity to changes to their total depletion. Therefore, we estimated the maximum sensitivity of near-extinction resources by selecting the binned sensitivities with $0.7 < C_\alpha < 1.2$ (limits were chosen *ad-hoc*), found peaks in these sensitivities using *scipy.signal.find_peaks*, then selecting the maximum peak value. Hyper-parameters: the threshold value at which a peak was detected *threshold* ad-hoc set to 0.01.

#### D.6 Numerically solving the self-consistency equations

The self-consistency equations describing the consumer and resource abundance distributions — eq.s (28) - (35) — cannot be solved analytically. Instead we needed to solve them numerically for each set of model parameter distributions. Please see `https://github.com/JamilaRowlandChandler/CRM-Resource-supply-vs-Stability` to see how this numerical routine is run.

To obtain solutions to our self-consistency equations, we used `NMinimize` using the `RandomSearch` method to run a non-linear least-squares minimisation. This routine traverses a large area of parameter space and does not assume the problem is convex, which helped us obtain a global solution. This routine solved the self-consistency equations from any initial conditions within the bounds of said equations (e.g., $\langle v_N \rangle < 0$, etc.), and could even solve in the infeasible region of the model.

The loss function $\mathcal{L}(\mathrm{SCE})$ minimised was the sum of the squared difference between the estimated value of each self-consistency equation and the value calculated from the self-consistency equations listed in (**??**).

$$\mathcal{L}(\mathrm{SCE}) = \sum_{p \in \mathrm{SCE}} (p_{i,\mathrm{estimated}} - p_{i,\mathrm{calculated}})^2, \text{ where SCE is the set } \left\{\phi_N, \langle N \rangle, \langle N^2 \rangle, \langle v_N \rangle\right\}. \tag{118}$$

We then numerically evaluated $\langle R \rangle, \langle R^2 \rangle, \langle \chi_R \rangle$ at the best solution.

##### D.6.1 Solving for the stability boundary

Firstly, we solved the self-consistency equations using the routine we just described. Then, we plugged the self-consistency equations into the expression describing how far the community was from the stability threshold, given by

$$\frac{1}{4\rho^2}\left[1 - \frac{2}{\left[\left[\mu_b + \mu_c\gamma^{-1}\langle N\rangle\right]^2 - 4\sigma_c\sigma_g\rho\gamma^{-1}\langle v_N\rangle\mu_I\right]^{\frac{1}{2}}}\left[\mu_b + \mu_c\gamma^{-1}\langle N\rangle - \frac{3\left[\mu_b + \mu_c\gamma^{-1}\langle N\rangle\right]\left[\sigma_c^2\gamma^{-1}\langle N^2\rangle + \sigma_b^2\right]}{2\left[\left[\mu_b + \mu_c\gamma^{-1}\langle N\rangle\right]^2 - 4\sigma_c\sigma_g\rho\gamma^{-1}\langle v_N\rangle\mu_I\right]}\right]\right.$$
$$+ \frac{1}{\left[\mu_b + \mu_c\gamma^{-1}\langle N\rangle\right]^2 - 4\sigma_c\sigma_g\rho\gamma^{-1}\langle v_N\rangle\mu_I}\left[\left[\mu_b + \mu_c\gamma^{-1}\langle N\rangle\right]^2 + \sigma_c^2\gamma^{-1}\langle N^2\rangle + \sigma_b^2 - \frac{5\left[\mu_b + \mu_c\gamma^{-1}\langle N\rangle\right]^2\left[\sigma_c^2\gamma^{-1}\langle N^2\rangle + \sigma_b^2\right]}{\left[\mu_b + \mu_c\gamma^{-1}\langle N\rangle\right]^2 - 4\sigma_c\sigma_g\rho\gamma^{-1}\langle v_N\rangle\mu_I} - \frac{3\left[\sigma_c^2\gamma^{-1}\langle N^2\rangle + \sigma_b^2\right]^2}{\left[\mu_b + \mu_c\gamma^{-1}\langle N\rangle\right]^2 - 4\sigma_c\sigma_g\rho\gamma^{-1}\langle v_N\rangle\mu_I}\right]\left.\vphantom{\frac{1}{1}}\right]$$
$$- \phi_N\gamma^{-1}. \tag{119}$$

When this expression equals 0, the community is on the stability threshold. For each value of $\rho$, we fitted a curve that was a function of $\sigma$ to the stability distance using scipy.optimize.curve_fit. We then interpolated the curve and found the value of $\sigma$ where the stability distance was closest to 0.

### E Extended information

#### E.1 The self-renewing and externally-supplied model can become equivalent when resource dynamics are neglected

Some studies have claimed that resource supply dynamics do not alter the phase transitions of the consumer-resource model [4, 5]. To understand why these models might be expected to exhibit equivalent phase transitions, we consider the limit where resource dynamics are fast relative to consumer-dynamics. Resources then remain at pseudo-steady state ($\mathrm{d}R_\alpha/\mathrm{d}t = 0$), allowing their abundances to be expressed in terms of consumers and substituted into the consumer dynamics to yield a consumer-only model:

$$\frac{\mathrm{d}N_i}{\mathrm{d}t} = N_i\left[r_i - \sum_j^S A_{ij}N_j\right] \tag{120}$$

| | Self-renewing | Externally-supplied |
|---|---|---|
| $r_i$ | $\sum_\alpha^M K_\alpha b_\alpha g_{i\alpha}\Theta(R_\alpha) - d_i$ | $\sum_\alpha^M [I_\alpha/b_\alpha]g_{i\alpha} - d_i$ |
| $A_{ij}$ | $\sum_\alpha^M K_\alpha g_{i\alpha}c_{j\alpha}\Theta(R_\alpha)$ | $\sum_\alpha^M [I_\alpha/b_\alpha^2]g_{i\alpha}c_{j\alpha}$ |

Here, $r_i$ represents the growth rate of each consumer in the absence of others, while $A_{ij}$ represents effective pairwise interspecies interaction strengths. For self-renewing resources, the Heaviside function $\Theta(R_\alpha)$ is 0 if resource $\alpha$ has gone extinct, and 1 otherwise. Externally-supplied resources cannot go extinct, consumer dynamics only reduce to eq. (120) assuming $b_\alpha \gg \sum_i^S c_{i\alpha}N_i$. These consumer-only models become nearly identical when $I_\alpha = b_\alpha = K_\alpha = 1$, but only in the fast-resource limit and for externally supplied resources, total consumption must be far smaller than outflux. In general, outside of this limit, these two models could show qualitatively distinct phase transitions.

#### E.2 The stability condition for externally-supplied vs self-renewing resources diverge when resource susceptibilities are not all equal

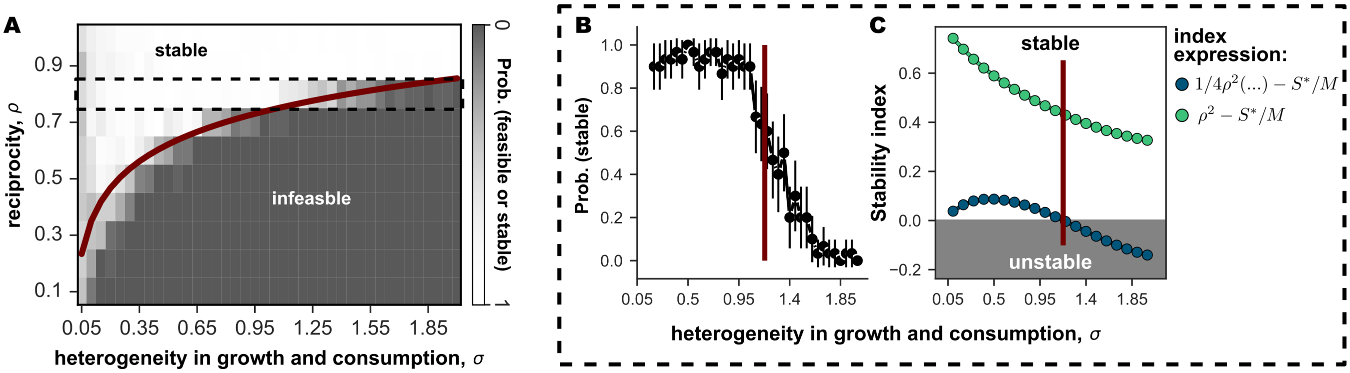

Figure S1: **Ecosystems with externally-supplied resources have a different stability condition to self-renewing resources** (Simulation parameters are given in Appendix E.) **(A)** Phase diagram of consumer-resource interaction reciprocity ($\rho$) and heterogeneity $\sigma$ for the model with externally-supplied resources. Each cell shows results from 30 simulated communities: dark grey shades indicate where simulations show unstable dynamics (maximum Lyapunov exponent $> 0$) which also turn out to be infeasible in this parameter range; and white indicates the communities are stable and feasible. The red line is the stability threshold, analytically derived from the externally-supplied model using the cavity method. **(B)** Simulations for $\rho = 0.8$, corresponding to the outlined area in panel (A). Error bars represent the 95% confidence interval, $n = 30$ communities. Red line = stability threshold for the externally-supplied model. **(C)** Stability indices derived under the assumption that resources have common susceptibilities (green) and with no assumptions (blue), both parametrised with the self-consistency equations solved for the externally-supplied model.

Here, we numerically solve the self-consistency equations for the externally-supplied model, and then estimate the stability boundary using eq. (63). The "full stability index" for the externally-supplied model is

$$\frac{1}{4\rho^2}\left[1 - \left\langle ... \right\rangle\right] - \frac{S^*}{M}, \tag{121}$$

and is $< 0$ when the community is stable and $= 0$ when the community reaches the stability threshold. Under the simple limit where all resources have the same susceptibilities $\chi_R$,

$$\rho^2 - \frac{S^*}{M^*}, \tag{122}$$

where $> 0$ when the community is stable and $= 0$ when the community reaches the stability threshold. The index in this limit is almost equivalent to the index for self-renewing resources (except resources can go extinct), but it is unclear whether the conditions remain the same or diverge when resource susceptibilities are not equal. To ensure the susceptibilities varied across resources, we varied the resource influx rate. Under this scenario, the full stability index captured the simulations well, but the simplified index could not: it predicted that the community should remain stable across (index $> 0$) the values of $\sigma_c$ tested, whereas the externally supplied condition correctly predicted a transition should occur (index $= 0$) (Fig. S1 C). This demonstrates that the stability condition for the externally-supplied model generally has a different form to the self-renewing condition.

#### E.3 Chaos emerges in the externally-supplied model under a narrow range of resource influx rates

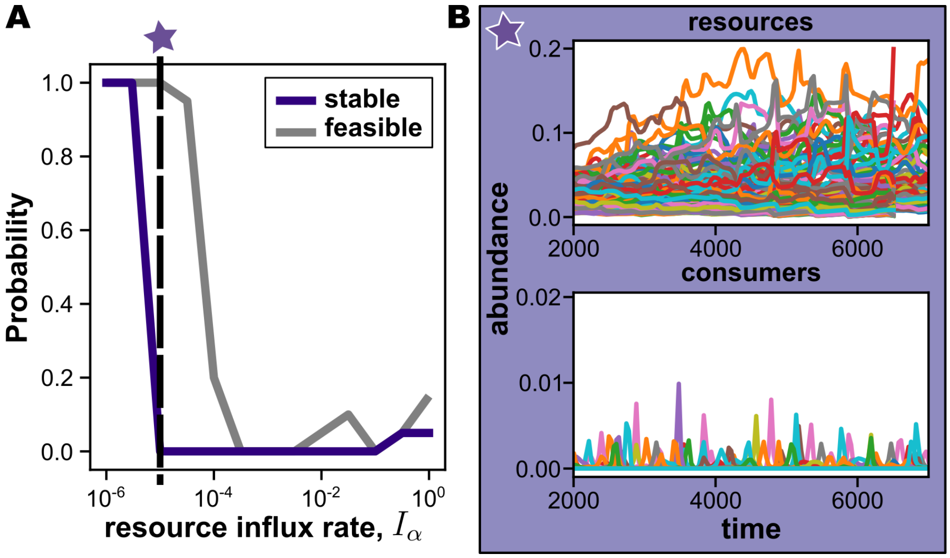

Figure S2: **Chaos in the consumer-resource model with externally-supplied resources** (Simulation parameters are given in Appendix E.) **(A)** Phase diagram of resource influx rate $I_\alpha$ (covaried with the dilution rate $b_\alpha$) for the model with externally-supplied resources. Each point shows results from 20 simulated communities, showing the proportion of communities with stable (purple) or feasible (grey) dynamics. Purple star indicates the critical resource influx/dilution rate where unstable and feasible dynamics i.e., chaos emerges. **(B)** Representative dynamics from the chaotic region of the externally-supplied model. Each curve in each plot represents a single consumer or resource, as labelled.
